# Genome resequencing, improvement of variant calling, and population genomic analyses provide insights into the seedlessness in the genus *Vitis*

**DOI:** 10.1101/863233

**Authors:** Myung-Shin Kim, Youn Young Hur, Ji Hong Kim, Soon-Chun Jeong

## Abstract

The seedlessness of grape derived from stenospermocarpy is one of the most prized traits of table or raisin grapes. It is controlled by a complex genetic system containing one major dominant gene and multiple minor recessive genes. Here, we collected dense variation data from high-depth resequencing data of seeded, seedless, and wild relative grape genomes sequenced to > 37x mean depth. Variant calls were made using a modified variant calling pipeline that was suitable for highly diverse interspecific grape accessions. The modified pipeline enabled us to call several million more variants than the commonly recommended pipeline. The quality was validated by Sanger sequencing data and subsequently supported by the genetic population structure and the phylogenetic tree constructed using the obtained variation data, results of which were generally consistent with known pedigree and taxonomic classifications. Variation data enabled us to confirm a major dominant gene and identify minor recessive loci for seedlessness. Incidentally, we found that grape cultivar Rizamat contains an ancestral chromosomal region of the major gene in Sultanina, a predominant seedlessness donor cultivar. Furthermore, we predicted new candidate causal genes including *Vitvi01g00455*, *Vitvi08g01528, and Vitvi18g01237* associated with the minor seedless-regulating loci, which showed high homology with genes that regulate seed development in *Arabidopsis*. This study provides fundamental insights relevant to variant calling from genome resequencing data of diverse interspecific hybrid germplasms such as grape and will accelerate future efforts aimed at crop improvement.

## Introduction

Grape is one of the most valuable fruit crops and is annually produced from 7.9 million ha globally (Faostat 2018). While most is processed into wine, a significant proportion (∼ 30%) is also destined for fresh consumption (table grape), dried into raisins, or processed into juice. However, ∼ 90% of grape produced from ∼ 14,000 ha in Korea is consumed as table grape (http://www.krei.re.kr/). Seedlessness is one of the most prized traits of table or raisin grape. Most of the seedless table grape cultivars with known pedigrees are derived from the stenospermocarpic variety Sultanina, also known as Sultanine or Thompson Seedless (Stout 1936; Bouquet and Danglot 1996). The most widely accepted hypothesis proposed for the inheritance of Sultanina-derived stenospermocarpic seedlessness is that the expression of three independently inherited recessive genes is controlled by a dominant regulator gene (Bouquet and Danglot 1996). Molecular markers tightly linked to the major dominant locus have been subsequently found and the locus was later named seed development inhibitor (*SDI*) (Lahogue *et al*. 1998). Quantitative trait loci (QTL) mapping studies have confirmed the existence of this major locus responsible for between 50% and 90% of total phenotypic variance in this trait, depending on the mapping population and trait evaluation (Lahogue *et al*. 1998; Doligez *et al*. 2002; Mejía *et al*. 2007). Several minor-effect QTL that could be recessive or modifying genes have also been described in these reports. The *SDI* locus is located on chromosome 18 (Mejía *et al*. 2007). In this region, the MADS-box gene *AGAMOUS-LIKE 11* (*VviAGL11*) has been proposed as a candidate for *SDI* (Costantini *et al*. 2008; Mejía *et al*. 2011) because the homologous gene in *Arabidopsis* is involved in ovule and carpel development. The direct role of *VviAGL11* in seed morphogenesis has been confirmed by its ectopic expression in the *Arabidopsis SEEDSTICK* mutant (Malabarba *et al*. 2017). An arginine-to-leucine substitution in *VviAGL11* has been postulated to be the major cause of seedlessness in grapevine cultivars (Royo *et al*. 2018). Thus far, however, minor seedless-regulating QTL have not been further elucidated at the molecular level.

Grape was the first fruit species to have its genome completely sequenced (Jaillon *et al*. 2007). Despite the early availability of a reference genome of *Vitis vinifera* subsp. *vinifera* PN40024 (derived from seeded Pinot Noir and close to homozygosity after 6-9 rounds of selfing), grape genomics has lagged behind other major crop plants (Schreiber *et al*. 2018) likely because of its high heterozygosity and long generation time. An analysis of approximately one thousand grape accessions using Vitis9kSNP array has revealed that *Vitis vinifera* L. subsp. *vinifera*, a domesticated grape species, has maintained high levels of genetic diversity and rapid linkage disequilibrium (LD) decay due to introgression from local wild progenitor *Vitis vinifera* L. subsp. *sylvestris* (Gmel.) Hegi during domestication (Myles *et al*. 2011). Despite a complex network of close pedigree relationships among elite cultivars, first-degree relationships are rare between wine and table grapes and among grapes from geographically distant regions. Recent studies have explored high levels of genomic variation in a few important cultivars (Di Genova *et al*. 2014; Cardone *et al*. 2016; Xu *et al*. 2016). A population-level genomics study of grape has investigated the domestication history of grape using genome resequencing data from nine *sylvestris* and 18 *vinifera* individuals (Zhou *et al*. 2017). However, comprehensive genome resequencing data at an average depth of ∼15.5× have only recently been used to investigate the population genomics of grapes to explore grape features other than domestication during the course of this study (Liang *et al*. 2019).

With an interest in elucidating seedless mechanisms in grape using genome-wide variation data, we have sequenced a diverse group of grape accessions (Hur *et al*. 2019). In this study, we report analyses of high-depth resequencing data from 14 seeded, 17 seedless, and two wild grape genomes sequenced to a mean depth > 37×. Such depth is likely sufficient for calling heterozygous genotypes (Ajay *et al*. 2011). In particular, because many *Vitis* hybrid cultivars have been generated for various purposes such as disease resistance, environmental adaptation, and flavors and are already cultivated, we obtained resequencing data from several cultivars generated from crosses between *V. vinifera* and its wild relative species such as *Vitis labrusca* L. However, the high diversity of our sequenced grape accession required a modification of the recommended popular variant calling pipeline. Data were first used to examine evolutionary relationships between seeded and seedless grapes and determine patterns of population structure and the decay of LD in the seeded and seedless grapes. We then used variation data to understand patterns of nucleotide diversity and LD surrounding the *SDI* locus and identify the minor recessive loci underlying seedlessness. The results of this study will be of great value both to grape breeders who are striving to more effectively harness haplotype variation at seedless-regulating loci to develop superior seedless grape cultivars and to genome researchers in general.

## Materials and methods

### Sample preparation and sequencing

We collected leaf tissues from a total of 27 individuals consisting of nine seeded, 16 seedless, and two wild grape accessions. Of these, 26 were selected from a grape collection grown in an experimental field of the National Institute of Horticultural and Herbal Science, Wanju, Korea while one accession, Rizamat Gs, was selected from a nursery at Gyeongsan, Korea (Table 1). Four out of eight seeded grape accessions were interspecific hybrids between *V. vinifera* and wild grape species. Four of 16 seedless grape accessions were hybrids. Genomic DNA was extracted from each leaf sample using Qiagen DNeasy plant kit. Paired-end sequencing libraries were constructed with an insert size of 500 bp using TruSeq DNA PCR-Free kit (Illumina, San Diego, CA) according to Illumina library preparation protocols. Libraries were then sequenced using Illumina HiSeq 4000 platform with 2 × 151-bp paired reads to a target coverage of 40X. Raw sequencing data were deposited in the Short Read Archive at NCBI (BioProject PRJNA485199). We also used Illumina raw reads with >37 coverage depths for five other seeded and one seedless cultivars (Da Silva *et al*. 2013; Di Genova *et al*. 2014; Gambino *et al*. 2017; Mercenaro *et al*. 2017; Zhou *et al*. 2017) that were downloaded from the Short Read Archive at NCBI (Table 1).

**Table 1.**
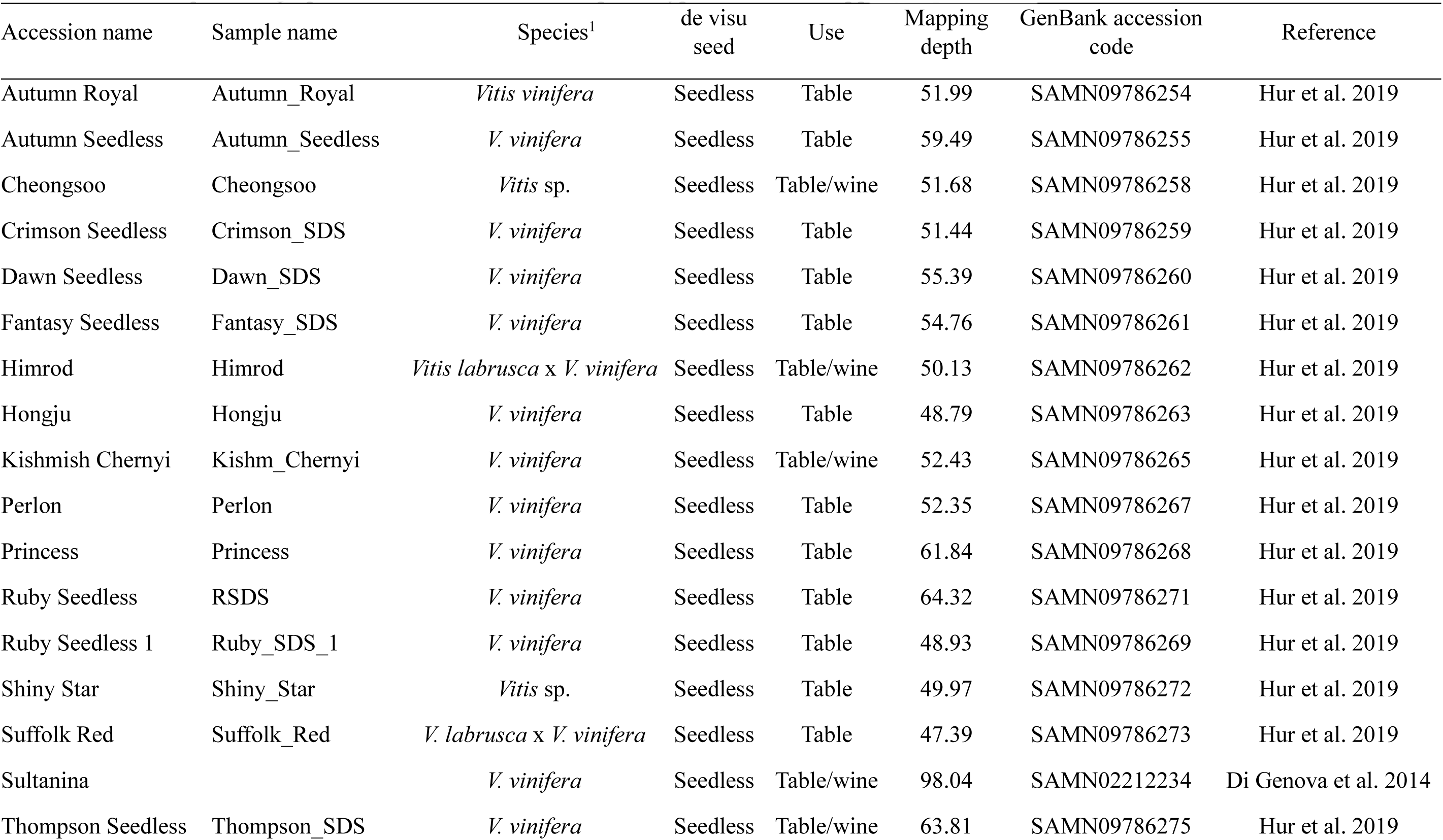
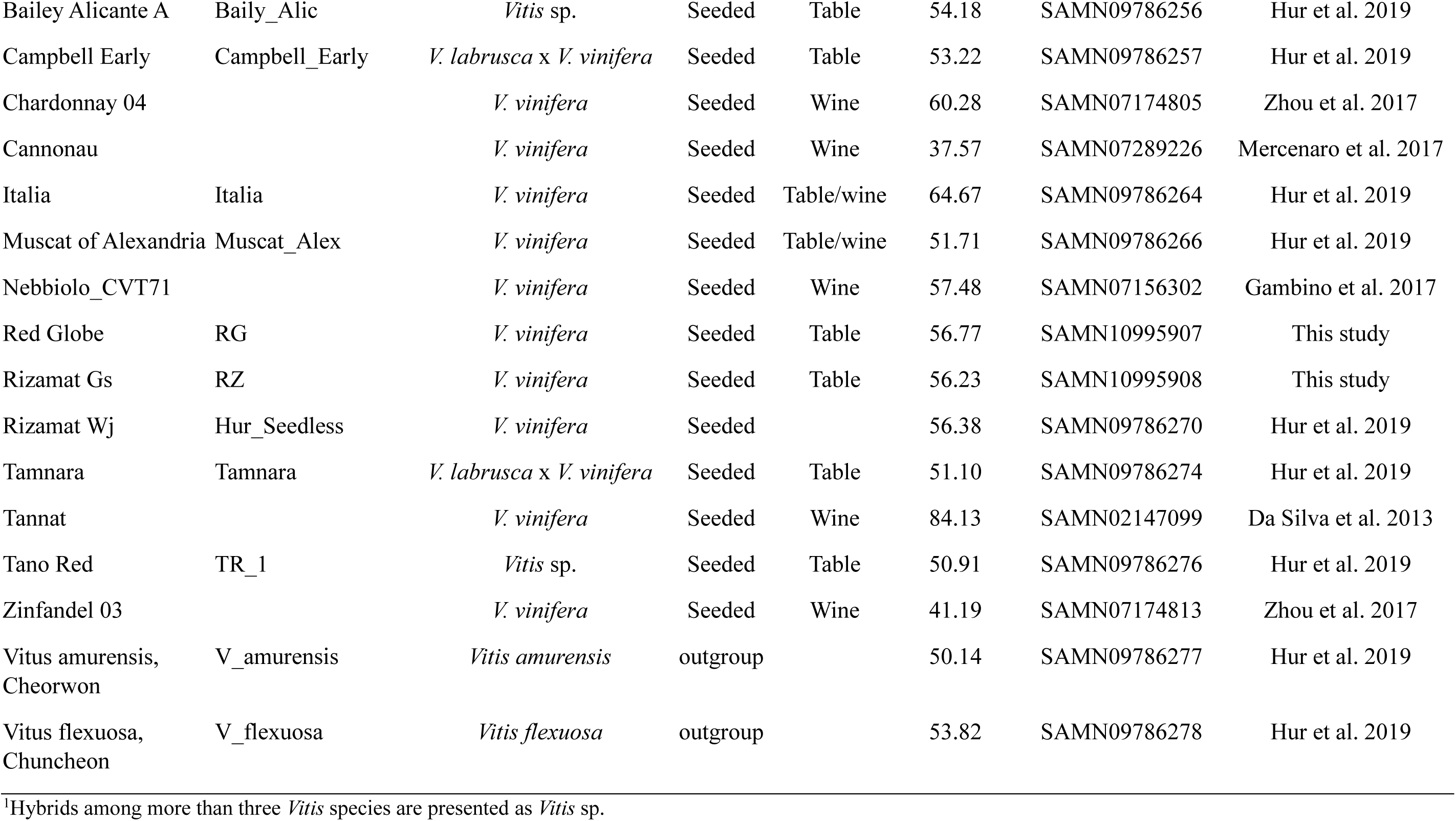
List of sequenced grape accessions and their seed phenotype and mean mapping depth

### Sequence alignment and variant calling

Short paired-end reads were quality checked using FastQC (http://www.bioinformatics.babraham.ac.uk/projects/fastqc/). We then essentially followed procedures described in the Genome Analysis Toolkit (GATK) Best Practices for data pre-processing (Depristo *et al*. 2011; Van Der Auwera *et al*. 2013) with some modifications. We used BWA (version 0.1.12) with default parameters (Li and Durbin 2009) to map genomic reads from each accession against *V. vinifera* Pinot Noir PN40024 reference genome (12X.v2) (Canaguier *et al*. 2017). Alignments were further checked for PCR duplicates using Picard (version 1.134) (http://picard.sourceforge.net/). We performed data pre-processing, including sorting operation and base recalibration, using GATK (version 4.0.1.2). A total of 457,245 known variants of grape genomes used in this study were release-40 data downloaded from https://plants.ensembl.org/index.html (accessed on 10 September 2018). Their coordinates were converted to 12X.v2 coordinates using python script provided by (Canaguier *et al*. 2017) before use. For variant calling, we used GATK (v. 3.8-0) UnifiedGenotyper available in GATK3 because our initial attempt using HaplotypeCaller implemented in GATK4 produced a much smaller number of SNPs from hybrids than from *V. vinifera*. After further processing by applying IndelRealigner, we called SNPs and indels with UnifiedGenotyper. Raw variant calling data were divided into SNPs and indels with SelectVariants function of GATK (v. 4.0.1.2). Hard-filtering was then performed for these raw SNP calls using VariantFiltration function of GATK (v. 4.0.1.2) according to the following threshold criteria: MappingQualityRankSum of < -12.5, polymorphism confidence scores (QUAL) < 30, genotype call quality divided by depth (QD) < 3.0, Phred-scaled P value of Fisher exact test for strand (FS) > 30.0, mapping quality (MQ) < 30.0, total depth of coverage (DP) < 200, and genotype-filter-expression depth of coverage (DP) < 15. Bi-allelic variants were then selected using VCFtools (version 0.1.15) (Danecek *et al*. 2011). To exclude erroneous variants in repetitive regions, variants with high mapping depth (> 4X reads per sample, where X is the mapping depth) in each sample were filtered. Allele balance (AB) was calculated and variants with AB < 30 in heterozygous genotypes were filtered. Variants with AB > 30 in homozygous genotypes in interspecific hybrids and wild relative species were converted to heterozygous genotypes. SNPs with missing rate > 30 % were removed using VCFtools (version 0.1.15) (Danecek et al., 2011). Filtering of raw indel calls was performed according to the following threshold criteria: ReadPosRankSum of < -20.0, QUAL < 30, QD < 2.0, and FS > 200, and DP < 200. Bi-allelic variants were then retained.

From this analysis, a total of 17,453,275 filtered SNPs and 3,109,464 filtered indels were defined as candidate variants. To perform population analyses, we further filtered these candidate SNPs using VCFtools (version 0.1.15) (Danecek et al., 2011) according to the following criteria non-ref-ac 1 --maf 0.05 --max-missing 0.9. Finally, we retained 5,373,452 high-quality SNPs in the data set.

### Variant calling with BCFtools

For variant calling, BCFtools (v. 1.9) (https://samtools.github.io/bcftools/bcftools.html), a variant calling project split from SAMtools package (Li 2011) was used to conduct analysis for output files from the above-described IndelRealigner step of GATK with the following options: bcftools mpileup -Ou -a FORMAT/AD, FORMAT/DP; bcftools call -Ov -mv -f GQ. VariantAnnotator function of GATK (v. 3.8-0) was used to add the following additional annotations: QualByDepth, MappingQualityRankSumTest, and FisherStrand. SNPs were then filtered using VariantFiltration function of GATK (v. 4.0.1.2) with criteria described above.

### Validation of variants

We validated candidate SNPs and indels called from genome resequencing data by Sanger sequencing of genomic DNA fragments PCR-amplified from twelve genes in four *V. vinifera* (Autumn Royal, Honju, Italia, Muscat of Alexandria, Rizamat Gs), two interspecific hybrid (Campbell Early and Cheongsoo), and one wild relative (*V. amurensis*) accessions. Primer sets were designed to amplify sequences in the genomic region of these genes (Table S1). In particular, three primer sets that would amplify overlapping gene fragments for the assembly of three contigs were designed to amplify sequences in the up-and down-stream and in exon/intron regions of *VviAGL11* in order to sequence the whole gene. As a result, the contigs from the same haplotypes were identified by specific polymorphisms in overlapping sequences. PCR amplifications were performed with 20 ng of grape genomic DNA using TAKARA LA Taq (Cat No. RR002A) with annealing temperature of 53°C for most of the genes except *VviAGL11*. Because gene fragments amplified for *VviAGL11* are larger than 3 kbp, we performed PCR amplifications using TAKARA LA Taq (Cat No. RR002A) with annealing temperature of 65°C. Because these called variants contained an appreciable rate of heterozygous variants, a given PCR product was subcloned into a plasmid for sequencing. At least three different clones for each haplotype were then sequenced. Sequences of both ends of a PCR amplicon were also determined directly from the amplified products to predict copy number of the amplicon based on sequence profiles.

### Population structure and relatedness

Population groups were inferred using a Bayesian model-based clustering method, fastSTRUCTURE (version 1.0) (Raj *et al*. 2014). FastStructure was run on default settings on 34 grape accessions including the Pinot Noir reference genome. The number of subpopulations (*K*) ranged from 1 to 12. Python script ChooseK incorporated with the FastStructure package was used to choose the number of subpopulations that could maximize the marginal likelihood. Results were graphically represented using STRUCTUREPLOT v1 (Ramasamy *et al*. 2014) with the Structure plot ordered by Q-value and accession names included as individual labels. Phylogeny among grape genomes was assessed using the neighbor-joining and bootstrap method implemented in MEGA7 (Kumar *et al*. 2016). Neighbor-joining trees were generated using *p*-distance measurement, pairwise deletion treatment, and 1000 bootstrap replicates to assess branch support. Principal Component Analysis (PCA) was performed using SMARTPCA with default setting (Patterson *et al*. 2006). For downstream analyses, the Pinot Noir reference genome, duplicated samples, and two wild relative grapes were excluded (*V. vinifera* Rizamat Wj, Ruby Seedless 1, and Sultanina; *V. amurensis* and *V. flexuosa*).

### Linkage disequilibrium (LD)

A total of 28 grape accessions consisting of 20 *V. vinifera*, and eight hybrids were separated and filtered using VCFtools (version 0.1.15) (Danecek *et al*. 2011) with --keep option and the following criteria: --non-ref-ac 1 --maf 0.1 --max-missing 0.9. Un-anchored (chr00), mitochondrial, and plastid sequences were also removed with --not-chr option. LD analysis was performed and plotted using PopLDdecay software (v. 3.4.0) (Zhang *et al*. 2018). Average *r^2^* of each 100 bp block was plotted using Plot_OnePop.pl script implemented in PopLDdecay.

### Predicting variant functional impact with SIFT

To predict functional effects of variants, Sorting Intolerant From Tolerant 4G (SIFT 4G) software (Vaser *et al*. 2016) was used. To create a grape database, uniref90 (https://www.uniprot.org/, download date: Feb 9th, 2019) was used as reference protein set. Annotation of *Vitis vinifera* 12X.v2 was downloaded from URGI (https://urgi.versailles.inra.fr/files/Vini/Vitis%2012X.2%20annotations/Vitis_vinifera_gene_annotation_on_V2_20.gff3.zip). Gff3 format was converted to Ensembl GTF format. Grape SIFT4G database was constructed using SIFT4G_Create_Genomic_DB implemented in SIFT4G. Functional effects of variants in coding regions of 33 grapes were predicted using SIFT4G annotator with default option.

### Genome scanning for selective signals

Twenty-eight grapes consisting of 13 seeded and 15 seedless grape accessions after excluding three duplicated samples and two wild relative samples were used for detecting selective sweep regions and logistic association. Monomorphic, minor allele frequency < 5 %, and missing rate > 10 % markers were filtered using VCFtools (version 0.1.15) (Danecek *et al*. 2011). Missing variants were imputed using BEAGLE v4.0 (Browning and Browning 2007) with default option. We performed a genome scan using a composite likelihood approach (XP-CLR) (Chen *et al*. 2010) updated by Hufford *et al*. (2012). Evidence for selection of seedlessness across the genome was evaluated by comparing seeded versus seedless grape genomes. Individual SNPs were assigned at positions along a Ri parental map derived from crosses between *V. vinifera* cv. Riesling cl.49 and *V. vinifera* cv. Gewürztraminer cl.643 (Ri×Gw) downloaded from URGI (https://urgi.versailles.inra.fr/download/vitis/Genetic_maps_Vitis_12X_V2.zip) (Canaguier *et al*. 2017). Coordinates of 12X.v2 genome assembly were applied using python script provided by (Canaguier *et al*. 2017) to calculate genetic per physical distance between markers in the Ri map. XP-CLR was performed with the following criteria: -w1 0.0005 200 100 –p1 0.7. In other words, XP-CLR scores of 100 bp window were calculated for maximum 200 SNPs per 0.05 cM genetic window and markers with a correlation level > 0.7 were down-weighted.

Seeded and seedless traits were encoded as binary traits of 1 (control) and 2 (case), respectively. Case-control logistic mixed model association test was performed using GENESIS (R package version 2.14.0) (Conomos *et al*. 2019) with default logistic mixed model association parameters assessed by Shenstone *et al*. (2018). The Manhattan plots of XP-CLR scores and logistic association p-values were constructed using qqman (Turner 2018) in R package.

## Results

### Variant calling

We analyzed resequencing data collected from a total of 33 grape accessions consisting of 13 seeded, 18 seedless, and two wild relative species with > 52x genome coverage (raw data) for variant calling (Table 1). Of these 33, we have recently reported 28 genome resequencing data that included eight seeded, 18 seedless grape cultivars, and two wild relatives as outgroups resequenced for this study (Hur et al. 2019). For the present study, we resequenced two additional seeded grape cultivars and added data from three previously reported accessions (Da Silva *et al*. 2013; Gambino *et al*. 2017; Mercenaro *et al*. 2017).

*V. vinifera* divided into subspecies *vinifera* and its progenitor subspecies *sylvestris* was the only species of food grape until the end of the 19^th^ century. However, since the outbreak of phylloxera at the end of the 19th century, interspecific hybrids between *V. vinifera* and other interfertile *Vitis* species were extensively introduced for disease resistance, different flavors, or adaptation to geographic areas other than the Mediterranean region (This et al., 2006). Because we selected important cultivars that were supposed to be better adapted in the Korean peninsula for resequencing, we included four seeded and four seedless interspecific hybrids (Table 1). After removing duplicate mapped reads, the mapped mean depth was > 37x for all accessions. More than 92% of the reference genome was covered by more than one read and > 87% were covered by more than five reads for all accessions. Thus, the mapping rate for hybrids and wild relatives is even better than those reported when resequencing data of wild rice *Oryza rufipogon* were mapped against the rice reference genome sequence from *Oryza sativa sub. japonica* (Xu *et al*. 2012). However, when we conducted raw candidate SNP calling using HaplotypeCaller implemented in GATK4 in our initial analysis (Fig. 1a), we obtained approximately one million less SNPs from most of the hybrids and wild relatives than those from *V. vinifera* (Figure S1a). This stands in stark contrast to the simple assumption that distantly related accessions would have higher genetic variation than closely related accessions, when compared to the reference genome sequence. This phenomenon became worse after a VariantFiltration step (Figure S2a). HaplotypeCaller calls SNPs and indels simultaneously via local *de-novo* assembly of haplotypes in an active region where it remains to be candidate variant loci on the basis of reads mapping through data pre-processing steps (Van Der Auwera *et al*. 2013). When we examined local assembly results using Integrative Genomics Viewer (Thorvaldsdottir *et al*. 2013), we observed that large portions of mapped reads became inactive after the local assembly, especially for hybrid and wild relative accessions (Figure S3). Local assembly results are used to obtain likelihoods of alleles for each variant in GVCF output files. This difference likely played a significant role at the multi-sample variant calling stage run through GenotypeGVCFs. In results, when we examined distribution of depth of coverage (DP) values from our grape accessions in raw SNP calling data obtained through GenotypeGVCFs, we found that hybrid and wild relative species contained significantly higher levels (several fold higher) of < 15 DP values than *V. vinifera* (Fig. 1b). Thus, for variant calling, we opted to use UnifiedGenotyper available in an older version of GATK which is a position-based caller without local re-assembly. Distribution patterns of DP values in raw SNP calling data obtained through UnifiedGenotyper were similar among *V. vinifera*, hybrids, and wild relative species (Fig. 1c). Moreover, the numbers of SNPs from hybrids and wild relative species were higher than those from *V. vinifera* (Figure S1b and Figure S2b), as expected based on their phylogenetic relationships. We also attempted to use BCFtools for variant calling. However, the number of raw SNPs or the number of filtered SNPs in each grape accession was approximately ten-fold lower than that from UnifiedGenotyper or HaplotypeCaller (Figure S2c). Therefore, we did not use it further.

**Fig. 1.**
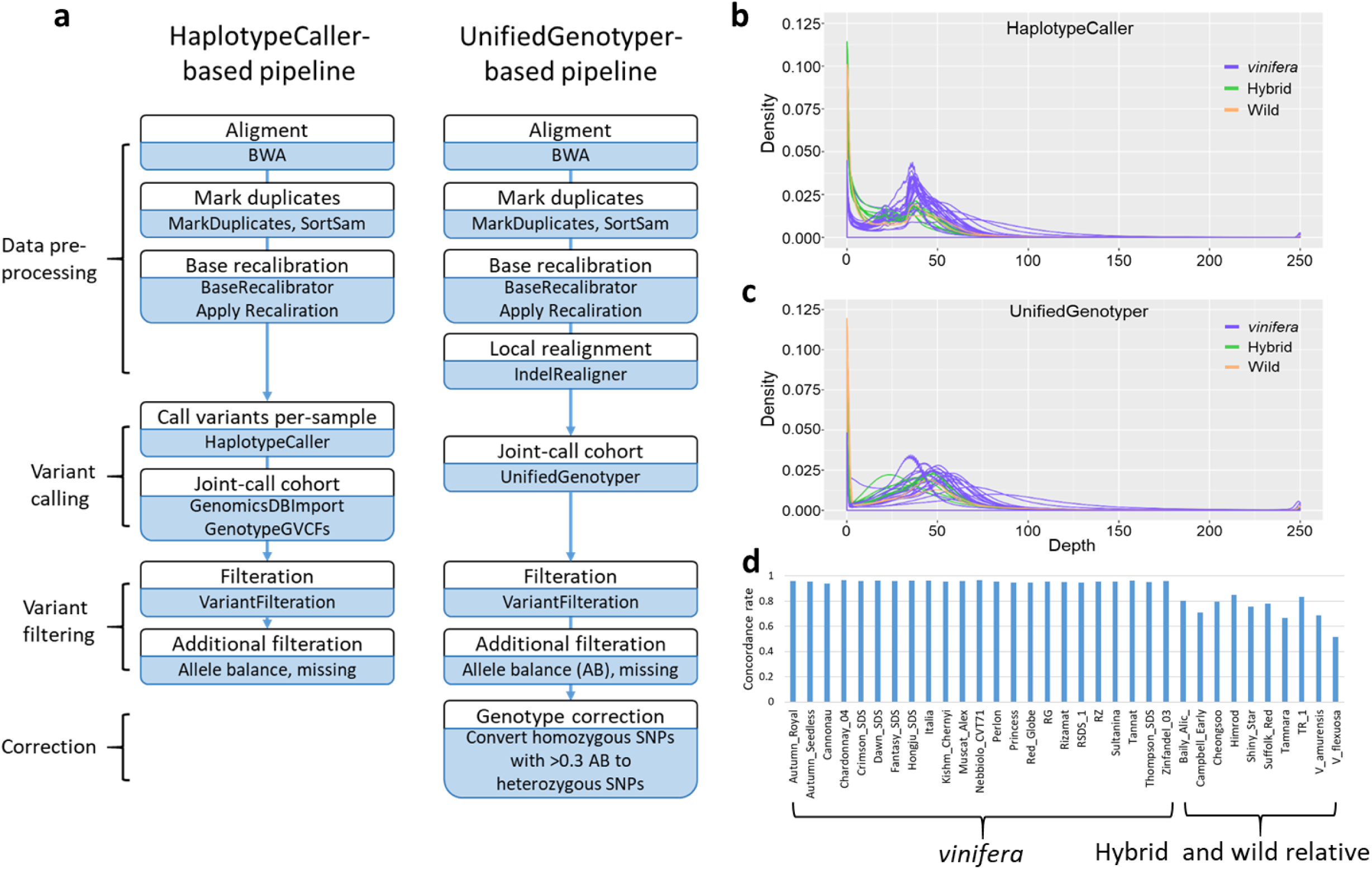
Comparison of workflows for variant discovery from grape genome sequencing data. **a** Workflows of HaplotypeCaller-based variant calling pipeline similar to the GATK Best Practices and UnifiedGenotyper-based variant calling pipeline. **b** Distribution of the depth of coverage (DP) per site from each of our grape accessions in raw SNP calling data obtained through GenotypeGVCFs using the HaplotypeCaller-based pipeline. *Vitis vinifera*, interspecific grape hybrids, and wild relative species are shown by green, blue, and orange lines. **c** Distribution of the DP per site from each of our grape accessions in raw SNP calling data obtained through UnifiedGenotyper using the UnifiedGenotyper-based pipeline. *Vitis vinifera*, interspecific grape hybrids, and wild relative species are shown by green, blue, and orange lines. Accessions with relatively higher DP per site are Cannonau (*V. vinifera*) and Suffolk Red (hybrid). **d** Genotype concordance rate of SNPs called by the GenotypeGVCFs with SNPs called by the UnifiedGenotyper in each of grape accessions. Only the SNPs that shared the grape reference genome coordinates were compared.

Differences in variant calling between HaplotypeCaller and UnifiedGenotyper might occur because some sequences from hybrids and wild relative species that are highly diverse relative to the grape reference genome sequence might have been treated as erroneous sequences. To examine this possibility, we compared *V. vinifera* variation data from only 23 *V. vinifera* and from all 33 grape accessions called through GenotypeGVCFs, and found no difference at identical sites between the two data sets. We also compared variant data called through GenotypeGVCFs with data called through UnifiedGenotyper from all 33 grape accessions. Interestingly, genotypes of SNPs that had identical coordinates between the two data sets were more than 93% similar in each V. *vinifera* accession and less than 85% similar in each hybrid and wild relative accession (Fig. 1d). These results indicate that UnifiedGenotyper might be more appropriate than HaplotypeCaller (recommended by developer) for variant calling analysis of distant relative species and their hybrids. It should be noted that application of UnifiedGenotyper for analysis of distant grape relative species, which are diploid, is an additional utility of this method because UnifiedGenotyper is recommended only for ploidy or pooled samples.

It was necessary to validate variant calls because the use of UnifiedGenotyper is not recommended by GATK Best Practices (Van Der Auwera *et al*. 2013). To validate variant calling results, we designed primer pairs from genomic regions of eleven randomly selected genes as well as of an *SDI*-encoding gene, *VviAGL11*, and performed Sanger sequencing. High-quality sequences determined by multiple clones in ten grape accessions ranged from 5.2 kb to 15.9 kb (Table S2). We also used *AGL11* sequences from Chardonnay and Sultanina reported by Malabarba *et al*. (2017). The sequences contained from 92 to 220 candidate SNP sites called through UnifiedGenotyper. Of the SNPs, 96.2% of UnifiedGenotyper SNP calls could be validated. However, only 77.4% of HaplotypeCaller SNP calls were validated (Table S2). As expected from the high similarity between UnifiedGenotyper and HaplotypeCaller SNP calls for *V. vinifera*, 96.2% of UnifiedGenotyper and 96.0% of HaplotypeCaller SNP calls were validated by Sanger sequences for the seven *V. vinifera*. However, 83.8% and 92.9% of UnifiedGenotyper and 38.6% and 55.8% of HaplotypeCaller SNP calls were validated by Sanger sequences for two hybrids and a *V. amurensis* accession, respectively. This large difference between the two pipelines were mainly due to missing calls of HaplotypeCaller. These findings likely explain the six million more SNPs from UnifiedGenotyper than that from HaplotypeCaller in total SNP calls.

When we examined the variant call format (VCF) file from UnifiedGenotyper, we found that many sites with higher than 30% of allelic balance were called homozygous SNPs. Interestingly, total numbers of SNP sites that were supposed to be erroneously called based on allelic balance were several hundred thousand for each of the hybrid and wild relative species accessions, with the highest number of 1.3 million seen for *V. flexuosa* (Table S3). However, these SNP sites numbered less than 10,000 for each *V. vinifera* accession, and these may be considered basal level errors. This phenomenon might have occurred because the UnifiedGenotyper caller purged alleles from more diverse reads of two haplotypes. Thus, we decided to convert these homozygous SNP to heterozygous SNPs. After this conversion, UnifiedGenotyper SNP calls for hybrid and wild relative accessions turned out to be highly accurate: 97.6% and 92.9% of UnifiedGenotyper SNP calls were validated by Sanger sequences for hybrids and wild relative accessions, respectively. In sum, we identified in all 33 accessions approximately 17.45 million candidate SNPs that went through the quality control filtering described in the Materials and Methods section below (Table S4). To obtain high-quality SNPs for population analyses, we excluded SNPs with < 10% minor allele frequency (MAF) and > 10% missing rate, yielding a total of 5,373,452 high-quality SNPs (Table S5). Relative to other accessions, Cannonau and two wild relative accessions consistently showed higher number of missing sites in all filtering methods and steps (Figure S4). In the case of Cannonau, this might occur due to its lowest mapping depth. For the two wild relatives, it might reflect some chromosomal regions that were too diverse for short reads to map properly.

### Population structure

A set of 5,373,452 high-quality SNPs was used to examine the genetic population structure and relationships among these 34 grape accessions including the Pinot Noir reference genome. To analyze the population structure, fastSTRUCTURE program (Raj *et al*. 2014) was used to estimate individual ancestry and admixture proportions assuming that *K* populations existed based on a maximum-likelihood method. The estimated marginal likelihood value plot in this analysis clearly supported the presence of three clusters (Figure S5). These three groups corresponded well with pedigree information (Fig. 2a). *V. vinifera* accessions formed two groups. Interestingly, *V. vinifera* accessions were roughly divided into wine and table grapes. These results are consistent with those of previous studies (Myles *et al*. 2011; Emanuelli *et al*. 2013) showing that close pedigree relationships are rare between wine and table grapes. Interspecific hybrids represented by Campbell Early and Tamara formed an independent group together with two wild relative grape species. However, the other hybrids appeared to be admixtures between *V. vinifera* and wild relative species. These grouping results that are consistent with pedigree and taxonomy indicated overall accuracy of variant calling in the present study. However, these results indicated that the grouping pattern based on SNPs was not related to that of grapes based on seeded and seedless phenotypes. Principal component analysis (PCA) results using SMARTPCA with default setting (Patterson *et al*. 2006) were consistent with grouping from the fastSTRUCTURE analysis (Fig. 2b and Figure S5).

**Fig. 2.**
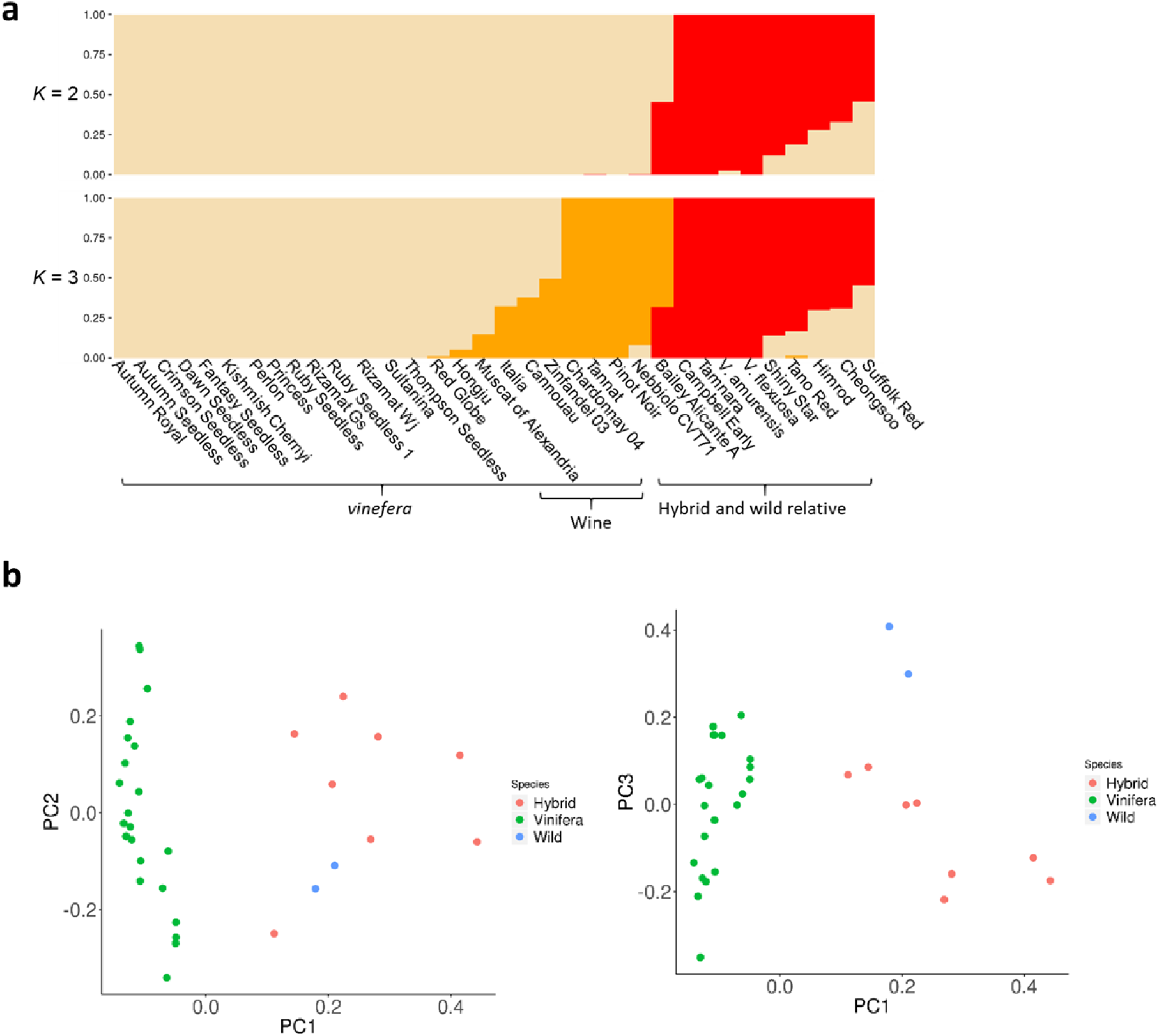
Grape population structure. **a** Population structure of 34 grape accessions estimated by fastSTRUCTURE. Each color represents one ancestral population. Each accession is represented by a vertical bar, and the length of colored segment in each vertical bar represents the proportion contributed by ancestral populations. **b** Principal components of SNP variation in grape accessions using whole-genome SNP data. The plots show the first three principal components. *Vitis vinifera*, interspecific hybrids, and wild relatives are shown by green, red, and blue dots, respectively.

The genetic population structure and relationships among these 34 grape accessions were further examined by constructing a neighbor-joining phylogenetic tree (Saitou and Nei 1987; Kumar *et al*. 2016). Consistent with fastSTRUCTURE results, these 34 grape accessions could be largely divided into three subclades (Fig. 3a). Group I contained only *V. vinifera* accessions except Bailey Alicante A and Sulfolk Red, while Group II consisted of all known interspecific hybrids. The two wild relative species formed an outgroup. In Group I, wine grapes clustered separately, as observed in the fastSTRUCTURE results. The grouping of Bailey Alicante A and Sulfolk Red with *V. vinifera* was consistent with results obtained from fastSTRUCTURE analysis showing that ancestry fraction from the *V. vinfera* group was considerably high in both accessions. Branch lengths among hybrids and wild relative species were longer than those among *V. vinifera* accessions, indicating that the diversity level of grapes obtained in this study was consistent with collection data in view of taxonomy and pedigree. It is notable that a tree constructed from SNP data without MAF filtration had much longer branch lengths for the two wild relative species than those in the tree with MAF filtration (Figure S6).

**Fig. 3.**
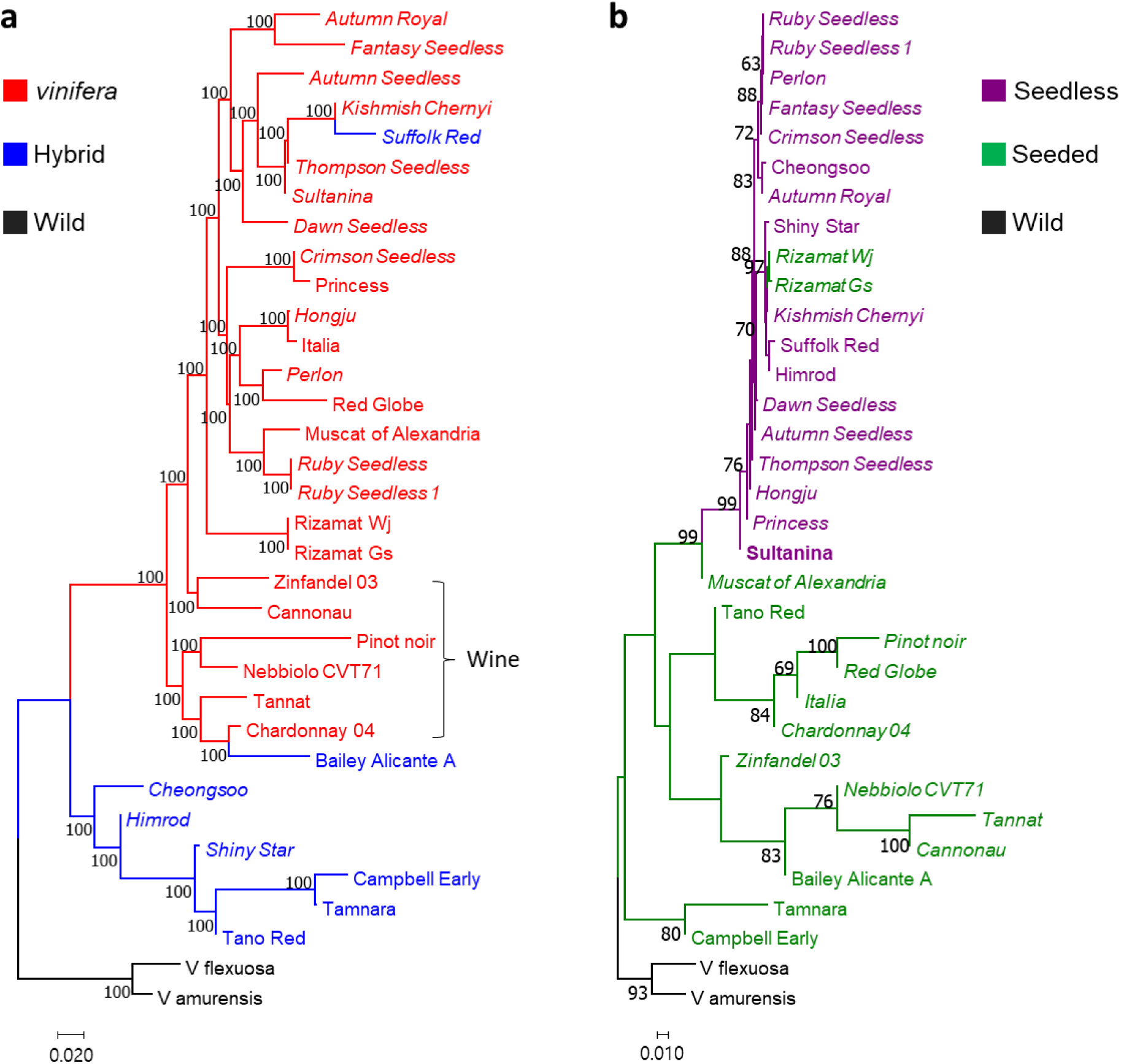
Phylogeny of grape. **a** Neighbor-joining phylogenetic tree of 34 grape nuclear genomes constructed using the 5,373,452 high-quality SNPs called from whole genome resequencing data. Accessions in the Neighbor-joining tree are represented by different colors: *Vitis vinifera* (red), interspecific hybrids (blue), and wild relatives (black). Group of wine grape accessions is indicated to emphasize their unique pedigree in this tree. Seedless accessions are in italic. **b** The tree constructed using 1,744 SNPs from 100-kb chromosomal region that contains the *SDI* locus in the central position. Percentages higher than 60 based on 1000 bootstrap replicates are shown above branches. Seeded grape accessions are represented by black letters and seedless accessions represented by red letters. *V. vinifera* accessions are in italic.

To estimate the LD patterns in different *V. vinifera* and interspecific hybrid groups, we calculated *r^2^* (Hill and Robertson 1968) between pairs of SNPs using PopLDdecay (Zhang *et al*. 2018). LD decayed to its half-maximum within approximately 11 kb for *V. vinifera* (Figure S7), which is similar to that for both wild and cultivated *V. vinifera* previously reported (Zhou *et al*. 2017). However, for interspecific hybrids, LD was high with a half-maximum of over 300 kb, a size that might be expected from a recently established population.

Although seeded and seedless grape accessions were intermixed within these two clearly separated subgroups in the phylogenetic tree constructed using genome-wide SNPs, seedless-regulating chromosomal regions might be confined within the diverse genetic background. To test this hypothesis, we constructed a tree using 1,744 SNPs from a 100 kb region surrounding the well-characterized *SDI* locus coding for *VviAGL11* (Fig. 3b). As expected, the tree formed two independent groups (seeded or seedless grape group) with the exception of Rizamat Wj and Rizamat Gs. Branch lengths within the seedless grape group appeared to be much shorter than those within the seeded grape group, supporting the notion of a single origin for the *SDI*-containing chromosomal region. To further examine *VviAGL11* sequences, we sequenced 8.9 kb of genomic DNA containing this gene. Although we had some difficulty due to preferential PCR amplification of parts of the haplotypes in several grape accessions (Figure S8), we were able to sequence several haplotypes of the full-length *VviAGL11* gene including a Sultanina mutant haplotype of Rizamat Gs. The Rizamat Gs *VviAGL11* sequence showed two SNPs in non-coding regions and only one SNP (arginine-to-leucine substitution site in *VviAGL11*) with several indels at the microsatellite repeat regions in non-coding regions. Moreover, the Rizamat Gs sequence grouped with seedless *VviAGL11* mutant sequences in our phylogenetic tree (Fig. 4). Most grape accessions contained two haplotypes of the *VviAGL11* gene on the basis of our phylogenetic tree constructed from upstream and first coding sequences of *VviAGL11* whose two haplotypes were PCR-amplified from accessions attempted in this study (Figure S9). However, among the approximately 300 SNPs and indels detected, only eight SNPs were located in coding regions, suggesting that coding regions have been well conserved. Of the eight, only two including the 197 arginine-to-leucine substitution site (Royo *et al*. 2018) were non-synonymous. The other non-synonymous SNP (210 threonine-to-alanine substitution) was interesting, however it is not likely to be another causal mutation because this site was not detected in our genome-wide logistic association scan described below. These results suggested that the *SDI*-containing region in our Rizamat clones might be an ancestral sequence where *SDI* mutation occurred in the Sultanina or its ancestor.

**Fig. 4.**
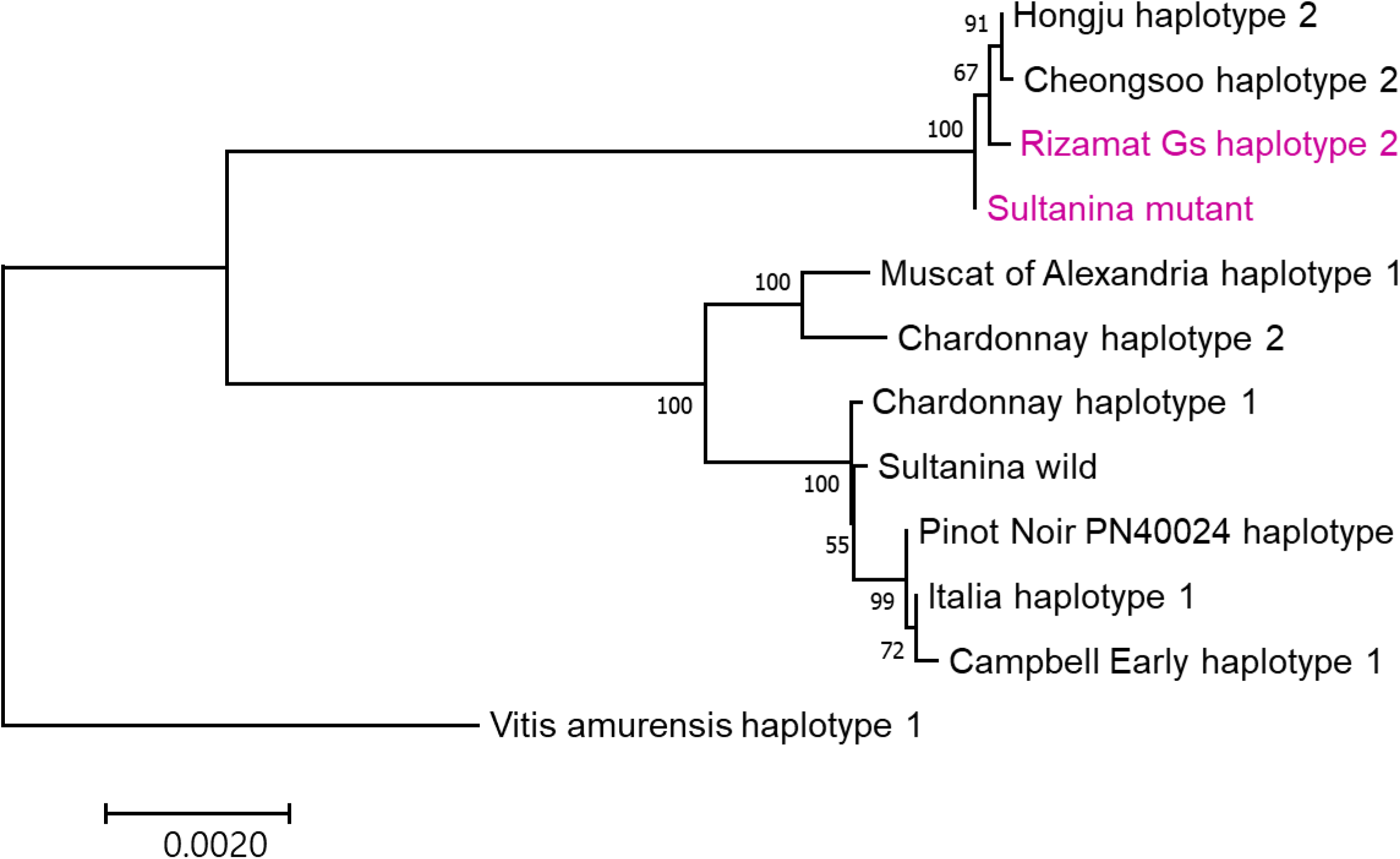
Neighbor-joining phylogenetic tree constructed from 12 haplotypes of 8.9 kb-genomic DNA sequences encoding the *VviAGL11* gene. Sultanina mutant and Rizamat Gs haplotype 2 are highlighted by purple letters.

We gathered three pairs of duplicated resequencing data. Ruby Seedless and Ruby Seedless 1 pair was generated due to a plant mislabeling. Rizamat clone pair (Rizamat Wj and Rizamat Gs) was obtained due to a problem with Rizamat Wj and a Thompson Seedless and Sultanina pair was generated after downloading Sultanina resequencing data that were publicly available. The main trunk of Rizamat Wj turned out to be dead and a shoot grew out from belowground during the 2018 growing season. Thus, we could not confirm the seedlessness phenotype of Rizamat Wj. Moreover, as Rizamat Wj grouped together with seedless accessions in the tree constructed from the *SDI*-containing chromosomal region, we opted to obtain resequencing data from another Rizamat clone, Rizamat Gs. The duplicate samples grouped together in our population structure and phylogeny analyses (Figs. 2 and 3). Their SNPs were approximately 99% similar to each other, assuring the high quality of our resequencing data. In the following analysis to identify seedless-regulating chromosomal regions, we excluded Ruby Seedless 1, Rizamat Wj, and Sultanina. Additionally, two wild relative species that were distantly grouped with other grape accessions were excluded. Finally, 13 seeded and 15 seedless grape accessions were analyzed. For analysis of this subset of the population, we used slightly lower number of high-quality SNPs due to exclusion of SNPs fixed in the subset.

### Identification of seedless-regulating chromosomal regions

Population structure and phylogenetic analyses showed that seeded and seedless grape accessions were intermixed within these two clearly separated subgroups. Such results indicate that seedless-regulating chromosomal regions that have undergone artificial selection might be localized within a diverse genetic background. Selective sweep regions most affected by artificial selection of seedlessness during grape breeding history likely correspond to the one major and three minor QTL predicted by genetic analysis (Bouquet and Danglot 1996). To test this hypothesis, we first used a likelihood method, the cross-population composite likelihood ratio XP-CLR (Chen *et al*. 2010) updated by Hufford *et al*. (2012), to scan for extreme allele frequency differentiation over extended linked regions (Fig. 5a). A total of 30 selective sweeps (Fig. 5a and Table S6) were detected in the highest 0.5% of XP-CLR values. Interestingly, one of the major peaks corresponded with the *SDI*-locus-residing chromosomal region, as a major dominant seedless-regulating QTL reported by numerous studies (Mejía *et al*. 2007; Mejía *et al*. 2011; Malabarba *et al*. 2017; Royo *et al*. 2018). As seedless grape genotypes at the major dominant gene are heterozygous or homozygous, a peak from this major locus relative to the recessive minor QTL chromosomal region might not be the highest. These results suggested that some selective sweeps detected might correspond to minor QTL regions where three independently inherited recessive genes controlled by the SDI locus reside.

**Fig. 5.**
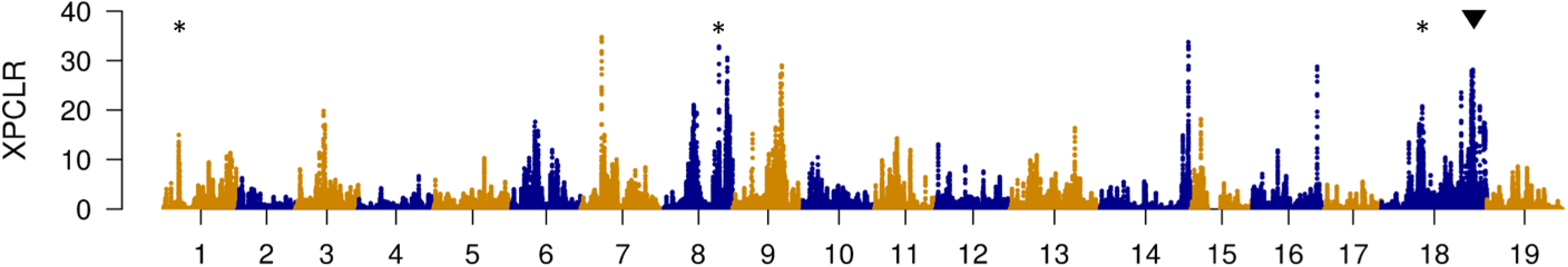
Genome-wide likelihood (XP-CLR) values for selection of seedlessness for seedless grape relatives to seeded grape accessions in 5-kb windows across the genome. The chromosome number is indicated along the x-axis. Chromosomal locations of *Vitvi01g00455*, *Vitvi08g01528, and Vitvi18g01237* associated with the minor seedless-regulating QTL and AGL11 associated with a major dominant QTL, which were predicted based on this XP-CLR analysis as well as logistic association and SIFT (Sorting Intolerant From Tolerant) analyses, are indicated by * and ▾, respectively.

Although our population size was only 28, consisting of 15 seedless and 13 seeded grape accessions, seedless-regulating SNPs might be more strongly associated with the seedlessness trait than other SNPs. Thus, we attempted to detect SNPs associated with seedlessness using a case–control logistic mixed model association test implemented in GENetic EStimation and Inference in Structured samples (GENESIS, R package version 2.14.0) software (Conomos *et al*. 2019), with correction of population structure that analyzed a binary phenotype of seeded or seedless phenotype. The highest peak correlated with the *SDI* locus on chromosome 18, unlike the XP-CLR analysis (Figure S10). Interestingly, an SNP with the highest -log10 *P* value of 5.051 in the highest peak was the arginine-to-leucine substitution site in *VviAGL11* identified as a causal mutation of the *SDI* locus. Comparison between XP-CLR and logistic association results showed that the majority of peaks were overlapping with each other. However, chromosome 19 contained two high peaks in logistic association scan while it did not contain a selective sweep and, vice versa, the end of chromosome 11 contained no significant peak in logistic association scan, however it did contain a high selective sweep peak. Those non-overlapping peaks between XP-CLR and logistic association scan results might be false positives generated by population structure or kinship. Thus, we focused on examining variants under the overlapping peaks in detail, with an expectation that we might pinpoint candidate causal genes for the postulated minor QTL.

### Assessment of variation patterns for causal gene prediction

Our SIFT (Sorting Intolerant From Tolerant) analysis (Vaser *et al*. 2016) predicted that 1,220 SNPs were deleterious in chromosomal regions of 50 kb to either side of the highest XP-CLR points in the 30 candidate selective sweeps detected (Table S6). Of the 1,220 SNPs, 41 SNPs in 34 genes showed -log_10_ *P* values higher than 2.5 from our logistic association (Table S7). In their milestone inheritance study, Bouquet and Danglot (1996) have shown that a system of three complementary recessive genes independently inherited is placed under the control of a completely dominant regulator gene *SDI*. When the *SDI* gene is heterozygous or homozygous dominant, expression of the seedless phenotype requires a minimum of two genes to be homozygous recessive. According to this model, most of the causal mutation sites at the *SDI* locus for seedless accessions should be homozygous or heterozygous non-reference alleles as Pinot Noir of the grape reference genome sequence accession is seeded, and most of causal mutations at the minor seedless-regulating QTL should be homozygous non-reference alleles.

We first examined variation patterns at a peak from 29.46 Mb to 30.46 Mb on chromosome 18, which includes the *SDI* locus (Fig. 5). Seven SNPs that were predicted to be deleterious using SIFT showed -log_10_ *P* values of higher than 2.5 from our logistic association scan (Table S7). Of the seven, genotype distribution of only one SNP in the seedless and seeded grape population was consistent with that predicted by the inheritance model of seedlessness. This SNP was heterozygous in all 15 seedless accessions tested, reference homozygous in 11 of 13 seeded accessions tested, and not called in 2 of the 13. Interestingly, it was the arginine-to-leucine substitution site identified as a causal mutation in *VviAGL11* encoding the *SDI* locus (Royo *et al*. 2018). Two of the remaining six SNPs were reference homozygous in one and two of 15 seedless accessions, respectively. One other was heterozygous in two seeded accessions, Italia and Nebbiolo_CVT71. Interestingly, all the remaining six were heterozygous in two Rizamat clones. The results are consistent with a tree constructed from a chromosomal region under this peak that showed clear separation of seeded and seedless grape groups with the exception of Rizamat Gs. Rizamat was developed by a cross of landraces in Uzbekistan close to Turkey where Sultanina was collected (http://www.vivc.de/) (Mirzaev and Djavacynce 2004). Thus, our SNP genotyping and the geographic origin of Rizamat suggest that this grape cultivar contained an ancestral chromosomal region of the *SDI* locus in Sultanina, a predominant seedlessness donor cultivar.

To examine variation patterns in the candidate selective sweeps of minor recessive seedless-regulating QTL, we classified all SNPs predicted to be deleterious into three groups using SIFT program in the candidate regions. Group I included SNPs that showed -log_10_ *P* values higher than 2.5 from our logistic association and were non-reference homozygous recessive in more than 10 of 15 seedless accessions tested. Group II included SNPs that showed -log_10_ *P* values higher than 2.5 from our logistic association and were non-reference homozygous recessive in less than 10 of 15 seedless accessions. Group III, which was excluded from further consideration, included SNPs that showed -log_10_ *P* values of lower than 2.5 from our logistic association test. Based on these criteria, 13 SNPs in four candidate selective sweeps were assigned to group I, whereas 21 SNPs in eight candidate selective sweeps were assigned to group II.

Six group I SNPs located within four genes were found at a peak from 4.3 Mb to 5.3 Mb on chromosome 1. Of the four genes, *Vitvi01g00455* was annotated as a cytosolic phosphoglucomutase (cPGM), a homolog of *Arabidopsis cPGM* (Egli *et al*. 2010) (Fig. 6a and Table S7). Loss of *cPGM* in *Arabidopsis* compromises male and female gametophyte development. Thus, *Vitvi01g00455* appears to be a strong candidate gene at this peak for a minor seedless-regulating QTL. A phylogenetic tree based on 1,669 SNPs from a 100-kb region surrounding *Vitvi01g00455* clearly separated seedless grape accessions with the non-reference homozygous recessive SNP and seedless and seeded grape accessions with the other genotypes (Fig. 6d). Only one group I SNP at the *Vitvi08g01528* gene model was found at a peak from 17.47 Mb to 28.47 Mb on chromosome 8 (Fig. 6b and Table S7). *Vitvi08g01528* was annotated as a basic helix-loop-helix transcription factor, a homolog of *Arabidopsis RETARDED GROWTH OF EMBRYO1* (*RGE1*) (Kondou *et al*. 2008). *Arabidopsis* RGE1 functions as a positive regulator in the endosperm at the heart stage of embryo development and exhibits pleiotropic phenotypes including small shriveled seeds and retardation of embryo growth. This annotation result suggests that *Vitvi08g01528* is the strongest candidate gene for a minor seedless-regulating QTL among the Group I SNPs. A phylogenetic tree based on 939 SNPs from a 100-kb region surrounding *Vitvi08g01528* clearly separated seedless grape accessions with the non-reference homozygous recessive SNP and seedless and seeded grape accessions with the other genotypes (Fig. 6e). Three group I SNPs at the *Vitvi08g02370* gene model were found at a peak from 20.20 Mb to 21.20 Mb on chromosome 8. This gene was annotated as a retrotransposon-related gene. Because this short gene with a coding region of 204 bp also has three deleterious SNPs, it is likely a pseudogene. On chromosome 18, the candidate peak was predicted to be from 12.83 Mb to 13.83 Mb. We assigned three SNPs to group I. They were mapped to three genes. Of the three genes, *Vitvi18g01230*, which is a short gene with a 183-bp coding region, was annotated as a retrotransposon-related gene. *Vitvi18g01245* was annotated as a retrotransposon-related probable LRR receptor-like serine/threonine-protein kinase with only one intron. *Vitvi18g01237* was annotated as a pentatricopeptide repeat protein, a homolog of *Arabidopsis MEF12*. *Arabidopsis MEF12* is involved in RNA editing in mitochondria (Hartel *et al*. 2013) and has not been studied for seed development. Considering the importance of RNA editing in plant development, *Vitvi18g01237* is a strong candidate for a minor seedless-regulating QTL. A phylogenetic tree based on 871 SNPs from 100-kb region surrounding *Vitvi18g01237* clearly separated seedless grape accessions with the non-reference homozygous recessive SNP and seedless and seeded grape accessions with the other genotypes (Fig. 6f). Taken together, the grouping pattern in three trees constructed from strong candidate minor QTL peaks is consistent with the inheritance model of seedlessness and indicates selection of this candidate selective sweep for seedlessness. This indicated that all three genes, which are known to be involved in seed development, are strong candidate causal genes for the minor seedless-regulating QTL.

**Fig. 6.**
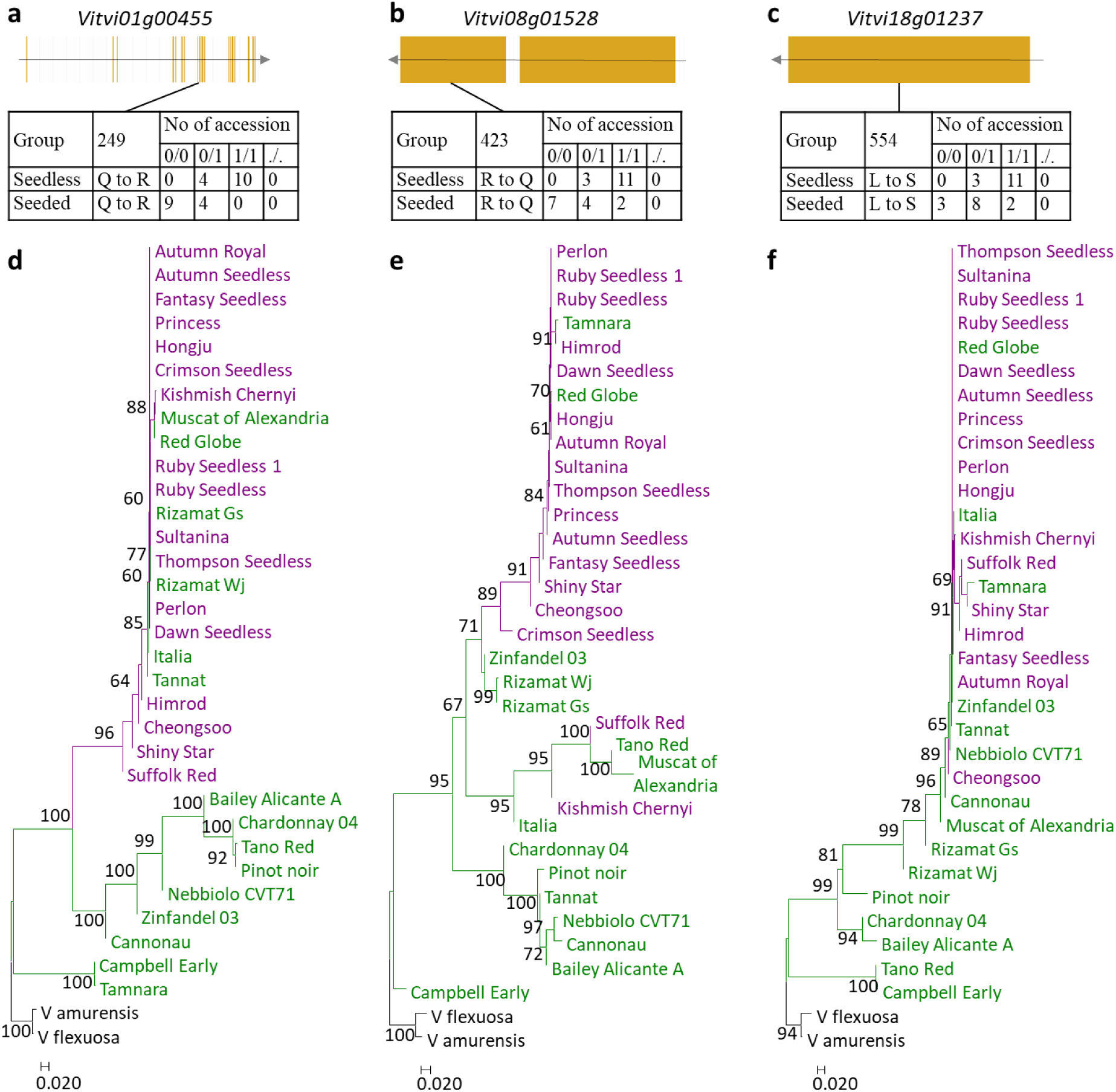
Genetic features for the candidate causal genes for minor recessive seedless-regulating candidate selective sweeps. Coding sequence structures of *Vitvi01g00455* (**a**), *Vitvi08g01528* (**b**), and *Vitvi01g01237* (**c**) and genotype distributions of candidate causal SNPs in these genes for 15 seedless and 13 seeded grape accessions tested. Reference homozygous genotype is indicated by 0/0, heterozygous 0/1, non-reference homozygous 1/1, and missing ./.. Amino acid positions are indicated by numbers. Neighbor-joining phylogenetic trees of 17 seedless and 14 seeded accessions constructed using 1,744 SNPs, 939 SNPs, and 871 SNPs from 100-kb chromosomal regions that contains the *Vitvi01g00455* (**d**), *Vitvi08g01528* (**e**), and *Vitvi01g01237* (**f**) genes in their central position, respectively. Percentages higher than 60 based on 1000 bootstrap replicates are shown above branches. Seeded grape accessions are in green, seedless accessions in purple, and two wild relatives in black.

Most of the group II SNPs appeared to be homozygous non-reference genotypes in less than five of 15 seedless accessions tested, thereby excluding the possibility of the SNPs-carrying genes for minor recessive seedless-regulating candidate genes. Five group II SNPs at five genes showed homozygous non-reference genotypes in seven or eight of 15 seedless accessions tested. Of the five genes, only *Vitvi08g01518* has been implicated in the process of seed development: its *Arabidopsis* homolog HD2B (At5g22650) functions as a genetic factor associated with seed dormancy. However, it is unlikely to be one of the minor recessive seedless-regulating QTL candidate genes because the strongest candidate *Vitvi08g01528* resides at the same candidate selective sweep. The 41 SNP sites selected tended to show homozygous non-reference genotype in none of 13 seeded accessions tested. Several SNPs were non-reference homozygous in only one or two of 13 seeded accessions tested. Interestingly, Rizamat Gs showed homozygous non-reference genotype in all six selected SNPs on chromosome 1. In the phylogenetic tree constructed from the chromosomal region (Fig. 6d), Rizamat clones grouped together with seedless grape accessions. The results support our notion that Rizamat has carried an ancestral form of the *SDI*-residing chromosomal region in Sultanina.

## Discussion

Development of variant calling methods from genome resequencing data have typically revolved around humans, livestock animals, and major crop plants, most of which contain diploid genomes. Numerous variant callers including UnifiedGenotyper and SAMtools have been proposed for variant calling of polyploidy species that are prevalent in the plant kingdom. Although none of these software packages is definitively recommended over others, many studies have successfully used the called variant data to address important biological questions (Clevenger *et al*. 2015). However, efforts to call variants in hybrids between relative species and in wild relative species that widely exist in woody species such as grape have been poorly done. Most grape resequencing data have been obtained from *V. vinifera*, which is the species used to obtain the reference genome sequence. Genome resequencing data of 472 *Vitis* accessions published while conducting this study included 108 wild relative grape species and 109 interspecific hybrid accessions. However, variants called using HaplotypeCaller were not validated by an experimental approach. In this study, we show that UnifiedGenotyper is better than HaplotypeCaller for SNP calling in hybrids and wild relative species. We also added an additional step to convert erroneously called homozygous SNPs to heterozygous SNPs. Therefore, our modified variant calling pipeline should provide insight for improvement of current variant callers to facilitate molecular genetic studies including marker-trait association studies for interspecific hybrids and wild relative species.

Several lines of evidence suggest that our predicted minor QTL are likely real. First of all, both XP-CLR and logistic association peaks with a shared high peak at the well-characterized *SDI* locus showed that the majority of peaks detected were overlapping. Trees constructed from peak-residing chromosomal regions tended to separate seeded and seedless grape accessions. Several of 30 peaks were located at the same physical locations as previously reported for minor QTL identified using seedless phenotypes in full-sibling F_1_ populations (Mejía *et al*. 2007; Costantini *et al*. 2008). Genotype distributions of deleterious substitutions in genes that reside at three overlapping peaks were supported by those proposed by Bouquet and Danglot (1996) in their milestone inheritance study of seedlessness. As results, we were able to suggest three strong candidate causal genes, namely *Vitvi01g00455*, *Vitvi08g01528, and Vitvi18g01237* as associated with the minor seedless-regulating QTL. It is difficult to pinpoint a causal gene underlying a minor QTL even with a large segregating population. In this study, based on analysis of high-density genome-wide SNP data, we have pinpointed several good candidate seedless-regulating genes that can be tested using techniques such as mutagenesis, transformation, and gene editing in the near future. This was made possible due to millions of high-quality genetic variants detected using our modified variant calling pipeline.

In this study, we have provided a large genome-wide variation dataset for seedless and seeded grape accessions with diverse genetic backgrounds. Because our initial variant calling efforts suggested that the current widely used variant calling pipeline had problems with interspecific hybrids and wild relative species, we modified the pipeline. Variation data from the modified pipeline were validated by Sanger sequencing. Our population structure and phylogenetic analysis using the resultant high-quality SNPs strongly supported known pedigree information as well as taxonomic grouping of these sequenced grape accessions, indicating that our modified pipeline was sound. The resulting millions of high-quality variations also provided an opportunity both to validate a major dominant seedless-regulating QTL and to predict minor recessive seedless-regulating QTL. Investigation of variation patterns at significant peaks allowed us to predict candidate causal genes that could regulate the seedless trait. Taken together, data generated in this study represent such a diverse grape genome background. They can now be used as dense markers of genome variation for marker-assisted mapping of important grape traits as well as for pinpointing agronomically important genes in grapes.

## Authors’ contributions

SCJ conceived the idea, performed data analysis, and wrote the manuscript. YYH conceived the project, performed grape selection for sequencing, and performed experiments. JHK performed sequencing of PCR products. MSK performed data analysis. All authors have read and approved the final version of the manuscript.

## Acknowledgements

We thank Dong Jun Im from National Institute of Horticultural and Herbal Science for assistance with collection of grape samples. This research was supported by a grant (PJ01242104) of the Rural Development Administration, Republic of Korea and by the Korea Research Institute of Bioscience and Biotechnology Research Initiative Program.

## Availability of data and materials

Short read data were deposited in the Short Read Archive at NCBI (BioProject PRJNA485199). Large datasets including SNP and indel calls and SIFT data are available from figshare repository (https://figshare.com/projects/Grape_resequencing_seedlessness_project/63569). The Sanger sequencing data from this study have been deposited with the GenBank data library under Accession Nos. MN243829–MN243907. A supplementary material file in the online of this article contains Figures S1 to S10 and Tables S1 to S7.

## References

1. Ajay, S. S., S. C. Parker, H. O. Abaan, K. V. Fajardo and E. H. Margulies, 2011 Accurate and comprehensive sequencing of personal genomes. Genome Res 21: 1498–1505.

2. Bouquet, A., and Y. Danglot, 1996 Inheritance of seedlessness in grape (*Vitis vinifera* L.). Vitis 35: 35–42.

3. Browning, S. R., and B. L. Browning, 2007 Rapid and accurate haplotype phasing and missing-data inference for whole-genome association studies by use of localized haplotype clustering. Am J Hum Genet 81: 1084–1097.

4. Canaguier, A., J. Grimplet, G. Di Gaspero, S. Scalabrin, E. Duchene et al., 2017 A new version of the grapevine reference genome assembly (12X.v2) and of its annotation (VCost.v3). Genom Data 14: 56–62.

5. Cardone, M. F., P. D’Addabbo, C. Alkan, C. Bergamini, C. R. Catacchio et al., 2016 Inter-varietal structural variation in grapevine genomes. Plant J 88: 648–661.

6. Chen, H., N. Patterson and D. Reich, 2010 Population differentiation as a test for selective sweeps. Genome Res 20: 393–402.

7. Clevenger, J., C. Chavarro, S. A. Pearl, P. Ozias-Akins and S. A. Jackson, 2015 Single Nucleotide Polymorphism Identification in Polyploids: A Review, Example, and Recommendations. Mol Plant 8: 831–846.

8. Conomos, M. P., S. M. Gogarten, L. Brown, H. Chen, K. Rice et al., 2019 GENESIS: GENetic EStimation and Inference in Structured samples (GENESIS): Statistical methods for analyzing genetic data from samples with population structure and/or relatedness, pp. in R package version 2.14.0.

9. Costantini, L., J. Battilana, F. Lamaj, G. Fanizza and M. S. Grando, 2008 Berry and phenology-related traits in grapevine (*Vitis vinifera* L.): from quantitative trait loci to underlying genes. BMC Plant Biol 8: 38.

10. Da Silva, C., G. Zamperin, A. Ferrarini, A. Minio, A. Dal Molin et al., 2013 The high polyphenol content of grapevine cultivar tannat berries is conferred primarily by genes that are not shared with the reference genome. Plant Cell 25: 4777–4788.

11. Danecek, P., A. Auton, G. Abecasis, C. A. Albers, E. Banks et al., 2011 The variant call format and VCFtools. Bioinformatics 27: 2156–2158.

12. DePristo, M. A., E. Banks, R. Poplin, K. V. Garimella, J. R. Maguire et al., 2011 A framework for variation discovery and genotyping using next-generation DNA sequencing data. Nat Genet 43: 491–498.

13. Di Genova, A., A. M. Almeida, C. Munoz-Espinoza, P. Vizoso, D. Travisany et al., 2014 Whole genome comparison between table and wine grapes reveals a comprehensive catalog of structural variants. BMC Plant Biol 14: 7.

14. Doligez, A., A. Bouquet, Y. Danglot, F. Lahogue, S. Riaz et al., 2002 Genetic mapping of grapevine (*Vitis vinifera* L.) applied to the detection of QTLs for seedlessness and berry weight. Theor Appl Genet 105: 780–795.

15. Egli, B., K. Kolling, C. Kohler, S. C. Zeeman and S. Streb, 2010 Loss of cytosolic phosphoglucomutase compromises gametophyte development in Arabidopsis. Plant Physiol 154: 1659–1671.

16. Emanuelli, F., S. Lorenzi, L. Grzeskowiak, V. Catalano, M. Stefanini et al., 2013 Genetic diversity and population structure assessed by SSR and SNP markers in a large germplasm collection of grape. BMC Plant Biol 13: 39.

17. FAOSTAT, 2018, pp.

18. Gambino, G., A. Dal Molin, P. Boccacci, A. Minio, W. Chitarra et al., 2017 Whole-genome sequencing and SNV genotyping of ‘Nebbiolo’ (Vitis vinifera L.) clones. Sci Rep 7: 17294.

19. Hartel, B., A. Zehrmann, D. Verbitskiy and M. Takenaka, 2013 The longest mitochondrial RNA editing PPR protein MEF12 in Arabidopsis thaliana requires the full-length E domain. RNA Biol 10: 1543–1548.

20. Hill, W. G., and A. Robertson, 1968 Linkage disequilibrium in finite populations. Theor Appl Genet 38: 226–231.

21. Hufford, M. B., X. Xu, J. van Heerwaarden, T. Pyhajarvi, J. M. Chia et al., 2012 Comparative population genomics of maize domestication and improvement. Nat Genet 44: 808–811.

22. Hur, Y. Y., M. S. Kim and S. C. Jeong, 2019 Towards identificationof haplotypes that control seedlessness of grape by genome resequencing. Acta Hort 1248: 171–178.

23. Jaillon, O., J. M. Aury, B. Noel, A. Policriti, C. Clepet et al., 2007 The grapevine genome sequence suggests ancestral hexaploidization in major angiosperm phyla. Nature 449: 463–467.

24. Kondou, Y., M. Nakazawa, M. Kawashima, T. Ichikawa, T. Yoshizumi et al., 2008 RETARDED GROWTH OF EMBRYO1, a new basic helix-loop-helix protein, expresses in endosperm to control embryo growth. Plant Physiol 147: 1924–1935.

25. Kumar, S., G. Stecher and K. Tamura, 2016 MEGA7: Molecular Evolutionary Genetics Analysis Version 7.0 for Bigger Datasets. Mol Biol Evol 33: 1870–1874.

26. Lahogue, F., P. This and A. Bouquet, 1998 Identification of a codominant scar marker linked to the seedlessness character in grapevine. Theor Appl Genet 97: 950–959.

27. Li, H., 2011 A statistical framework for SNP calling, mutation discovery, association mapping and population genetical parameter estimation from sequencing data. Bioinformatics 27: 2987–2993.

28. Li, H., and R. Durbin, 2009 Fast and accurate short read alignment with Burrows-Wheeler transform. Bioinformatics 25: 1754–1760.

29. Liang, Z., S. Duan, J. Sheng, S. Zhu, X. Ni et al., 2019 Whole-genome resequencing of 472 Vitis accessions for grapevine diversity and demographic history analyses. Nat Commun 10: 1190.

30. Malabarba, J., V. Buffon, J. E. A. Mariath, M. L. Gaeta, M. C. Dornelas et al., 2017 The MADS-box gene *Agamous-like 11* is essential for seed morphogenesis in grapevine. J Exp Bot 68: 1493–1506.

31. Mejía, N., M. Gebauer, L. Muñoz, N. Hewstone, C. Muñoz et al., 2007 Identification of QTLs for seedlessness, berry size, and ripening data in a seedless x seedless table grape progeny. Am J Enol Vitic 58: 499–507.

32. Mejía, N., B. Soto, M. Guerrero, X. Casanueva, C. Houel et al., 2011 Molecular, genetic and transcriptional evidence for a role of VvAGL11 in stenospermocarpic seedlessness in grapevine. BMC Plant Biol 11: 57.

33. Mercenaro, L., G. Nieddu, A. Porceddu, M. Pezzotti and S. Camiolo, 2017 Sequence polymorphisms and structural variations among four Grapevine (*Vitis vinifera* L.) cultivars representing Sardinian agriculture. Front Plant Sci 8: 1279.

34. Mirzaev, M. M., and U. M. Djavacynce, 2004 The Schroeder Institute in Uzbekistan: Breeding and germplasm collections. HortSci 39: 917–921.

35. Myles, S., A. R. Boyko, C. L. Owens, P. J. Brown, F. Grassi et al., 2011 Genetic structure and domestication history of the grape. Proc Natl Acad Sci U S A 108: 3530–3535.

36. Patterson, N., A. L. Price and D. Reich, 2006 Population structure and eigenanalysis. PLoS Genet 2: e190.

37. Raj, A., M. Stephens and J. K. Pritchard, 2014 fastSTRUCTURE: variational inference of population structure in large SNP data sets. Genetics 197: 573–589.

38. Ramasamy, R. K., S. Ramasamy, B. B. Bindroo and V. G. Naik, 2014 STRUCTURE PLOT: a program for drawing elegant STRUCTURE bar plots in user friendly interface. Springerplus 3: 431.

39. Royo, C., R. Torres-Perez, N. Mauri, N. Diestro, J. A. Cabezas et al., 2018 The Major Origin of Seedless Grapes Is Associated with a Missense Mutation in the MADS-Box Gene VviAGL11. Plant Physiol 177: 1234–1253.

40. Saitou, N., and M. Nei, 1987 The neighbor-joining method: a new method for reconstructing phylogenetic trees. Mol Biol Evol 4: 406–425.

41. Schreiber, M., N. Stein and M. Mascher, 2018 Genomic approaches for studying crop evolution. Genome Biol 19: 140.

42. Shenstone, E., J. Cooper, B. Rice, M. Bohn, T. M. Jamann et al., 2018 An assessment of the performance of the logistic mixed model for analyzing binary traits in maize and sorghum diversity panels. PLoS One 13: e0207752.

43. Stout, A. B., 1936 Seedlessness in grapes. N Y State Agricult Expt Stat Tech Bull (Geneva) 238.

44. Thorvaldsdottir, H., J. T. Robinson and J. P. Mesirov, 2013 Integrative Genomics Viewer (IGV): high-performance genomics data visualization and exploration. Brief Bioinform 14: 178–192.

45. Turner, S. D., 2018 qqman: an R package for visualizing GWAS results using Q-Q and manhattan plots. Journal of Open Source Software 3: 731.

46. Van der Auwera, G. A., M. O. Carneiro, C. Hartl, R. Poplin, G. Del Angel et al., 2013 From FastQ data to high confidence variant calls: the Genome Analysis Toolkit best practices pipeline. Curr Protoc Bioinformatics 43: 11 10 11–33.

47. Vaser, R., S. Adusumalli, S. N. Leng, M. Sikic and P. C. Ng, 2016 SIFT missense predictions for genomes. Nat Protoc 11: 1–9.

48. Xu, X., X. Liu, S. Ge, J. D. Jensen, F. Hu et al., 2012 Resequencing 50 accessions of cultivated and wild rice yields markers for identifying agronomically important genes. Nat Biotechnol 30: 105–111.

49. Xu, Y., Z. Gao, J. Tao, W. Jiang, S. Zhang et al., 2016 Genome-Wide Detection of SNP and SV Variations to Reveal Early Ripening-Related Genes in Grape. PLoS One 11: e0147749.

50. Zhang, C., S. S. Dong, J. Y. Xu, W. M. He and T. L. Yang, 2018 PopLDdecay: a fast and effective tool for linkage disequilibrium decay analysis based on variant call format files. Bioinformatics.

51. Zhou, Y., M. Massonnet, J. S. Sanjak, D. Cantu and B. S. Gaut, 2017 Evolutionary genomics of grape *(Vitis vinifera* ssp. *vinifera*) domestication. Proc Natl Acad Sci U S A 114: 11715–11720.

