## Supplementary_material_Figures_S1_to_S10_and_Tables_S1_to_S7 for "Genome resequencing, improvement of variant calling, and population genomic analyses provide insights into the seedlessness in the genus *Vitis*"

### Supplementary Figures S1 to S10 and Tables S1 to S7 for

### Genome resequencing, improvement of variant calling, and population genomic analyses provide insights into the seedlessness in the genus *Vitis*

**Kim et al., 2019**

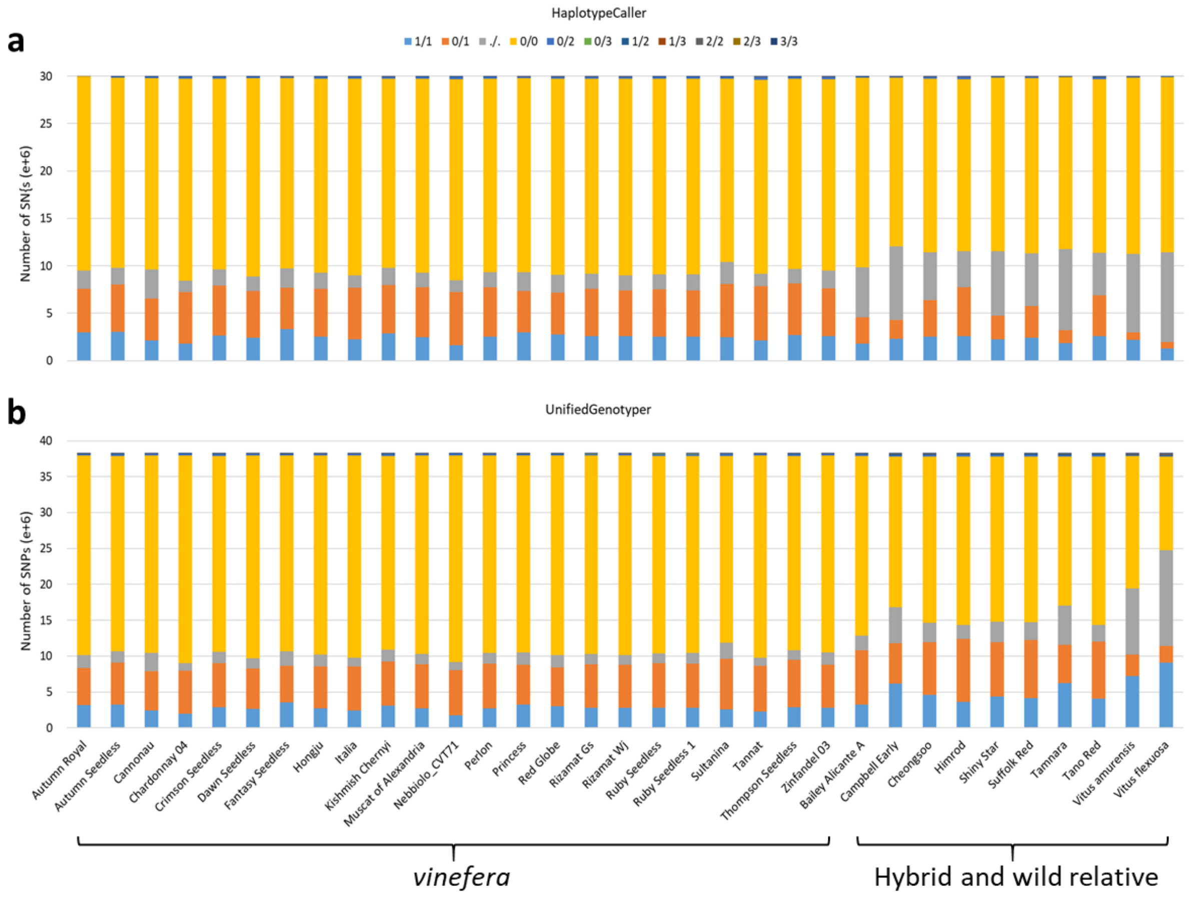


Figure S1. Distribution of SNP genotypes. **a** Distribution of 29.97 million raw candidate SNPs obtained using HaplotypeCaller for 33 grape accessions. **b** Distribution of 38.27 million raw candidate SNPs obtained through UnifiedGenotyper for 33 grape accessions. Reference allele is indicated by 0, non-reference alleles are 1, 2, and 3, and missing is indicated by a dot (.). Order of accessions and color scheme of SNP genotypes are the same in two panels.

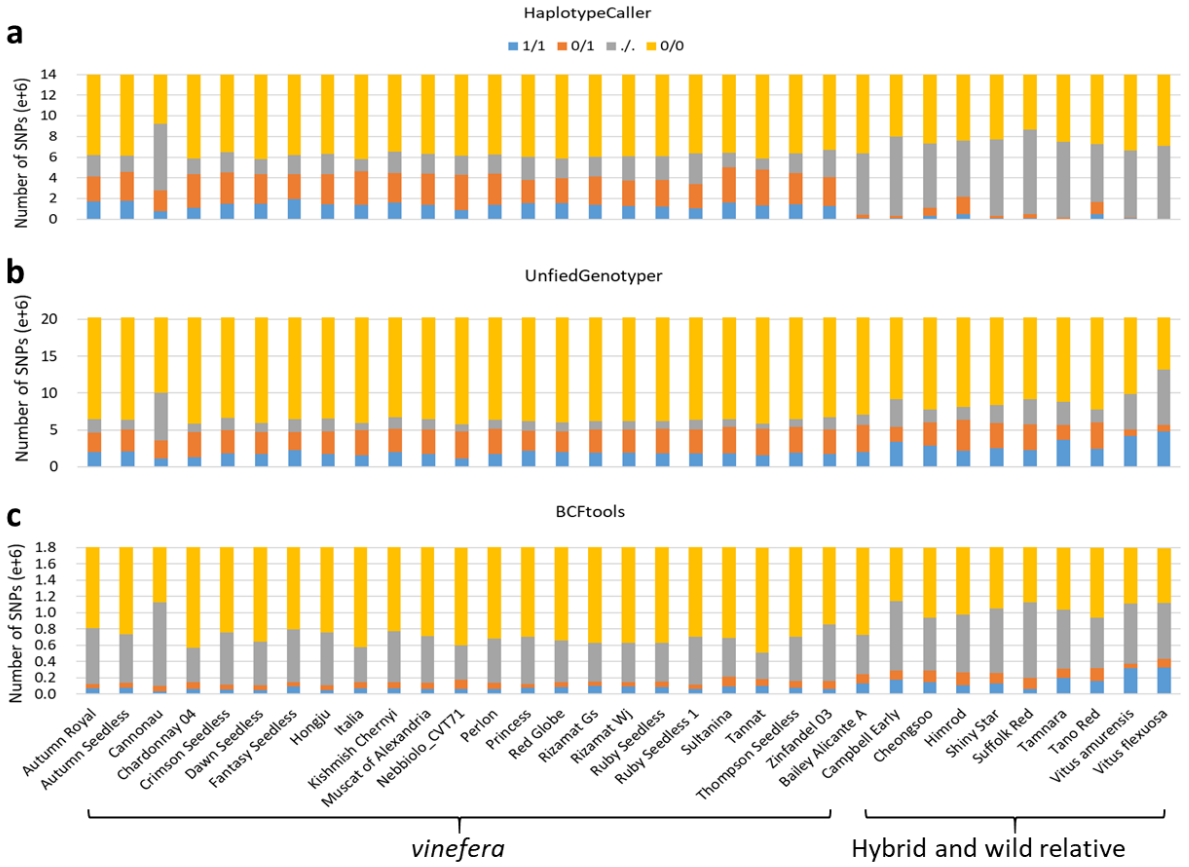


Figure S2. Distribution of SNP genotypes. **a** Distribution of 13.99 candidate bi-allelic SNPs remained after VariantFiltration filtering of raw candidate SNPs obtained using HaplotypeCaller for 33 grape accessions. **b** Distribution of 20.21 million candidate bi-allelic SNPs remained after VariantFiltration filtering of raw candidate SNPs obtained through UnifiedGenotyper for 33 grape accessions. **c** Distribution of 1.80 million candidate bi-allelic SNPs remained after VariantFiltration filtering of raw candidate SNPs obtained through BCFtools for 33 grape accessions. Reference allele is indicated by 0, non-reference alleles are 1, 2, and 3, and missing is indicated by a dot (.). Order of accessions and color scheme of SNP genotypes are the same in three panels.

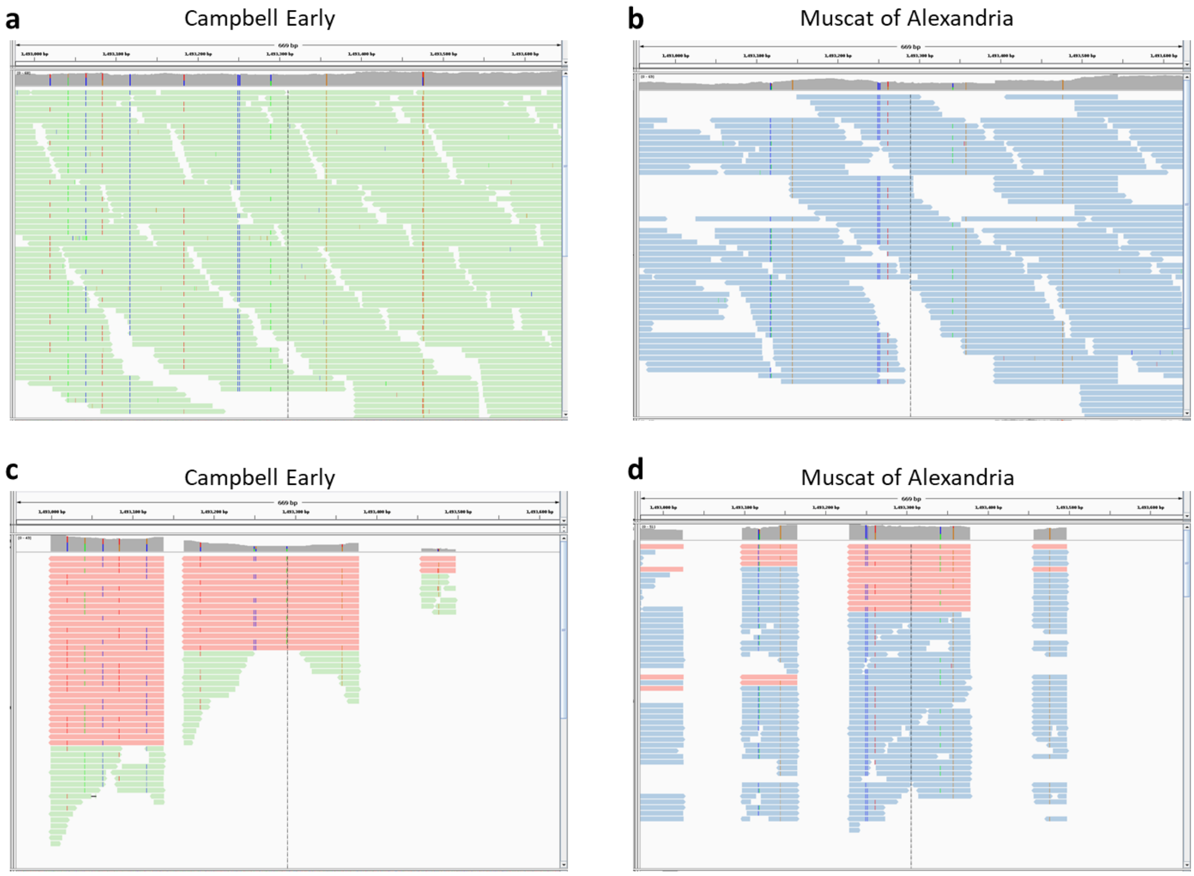


Figure S3. Representative read alignment views on Integrative Genomics Viewer (v. 2.3.10) (Thorvaldsdottir et al. 2013) to show production of inactive reads through HaplotypeCaller in the vicinity of 1,493,300 base position on chromosome 1. **a** Read alignment views in BAM data files after removing duplicate reads using MarkDuplicates in genome resequencing data of Campbell Early (panel **a**) and Muscat of Alexandria (panel **b**). Read alignment views in BAM data files obtained after local *de-novo* assembly using HaplotypeCaller in genome resequencing data of Campbell Early (panel **c**) and Muscat of Alexandria (panel **d**). Inactive reads are in red color. Campbell Early, a interspecific hybrid accession, showed higher portions of inactive reads than Muscat of Alexandria, a *Vitis vinifera* accession, at each SNP site.

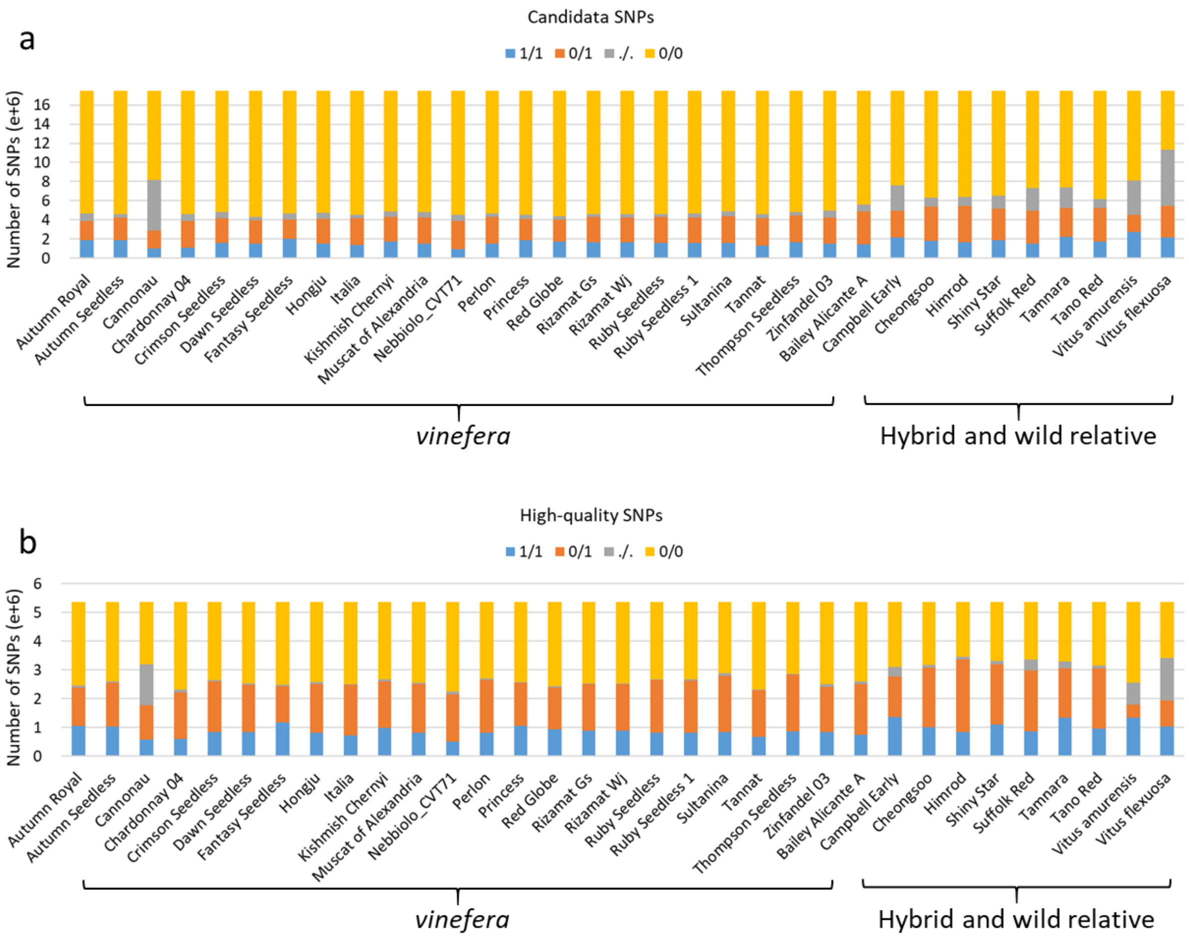


Figure S4. Distribution of SNP genotypes. **a** Distribution of 17.45 million candidate SNPs obtained using our UnifiedGenotyper-based pipeline for 33 grape accessions. **b** Distribution of 5,373,452 high-quality SNPs obtained through our UnifiedGenotyper-based pipeline for 33 grape accessions. Reference allele is indicated by 0, non-reference alleles are 1, 2, and 3, and missing is indicated by a dot (.). Color scheme of SNP genotypes is shown in legend above chart.

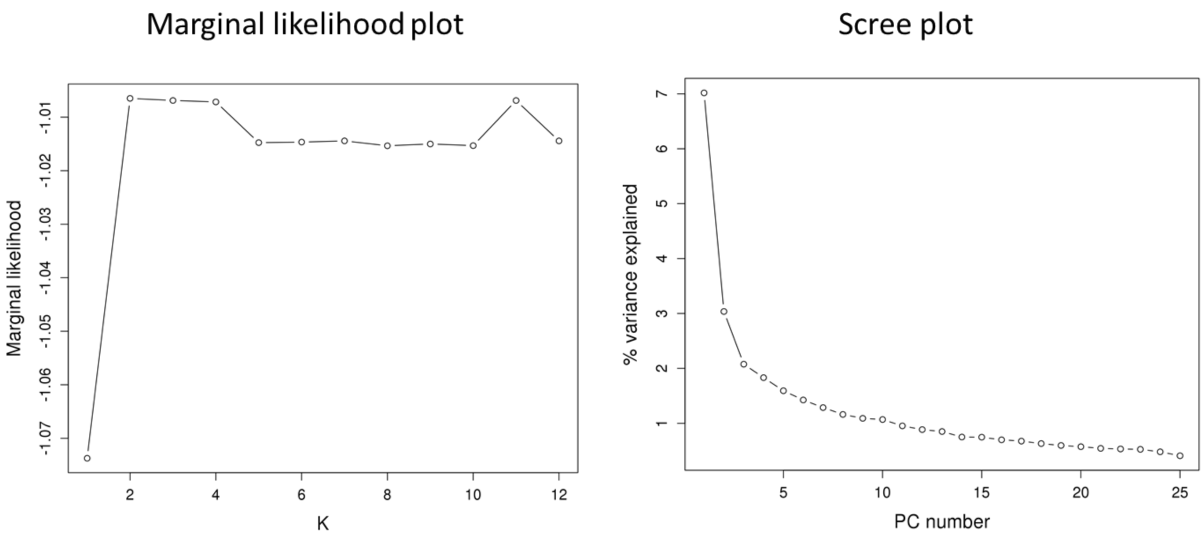


Figure S5. Plot of marginal likelihood values from fastSTRUCTURE analysis of the 34 grape accessions and scree plot of the principal component (PC) number and their contribution to variance from principal component analysis of the 34 grape accessions.

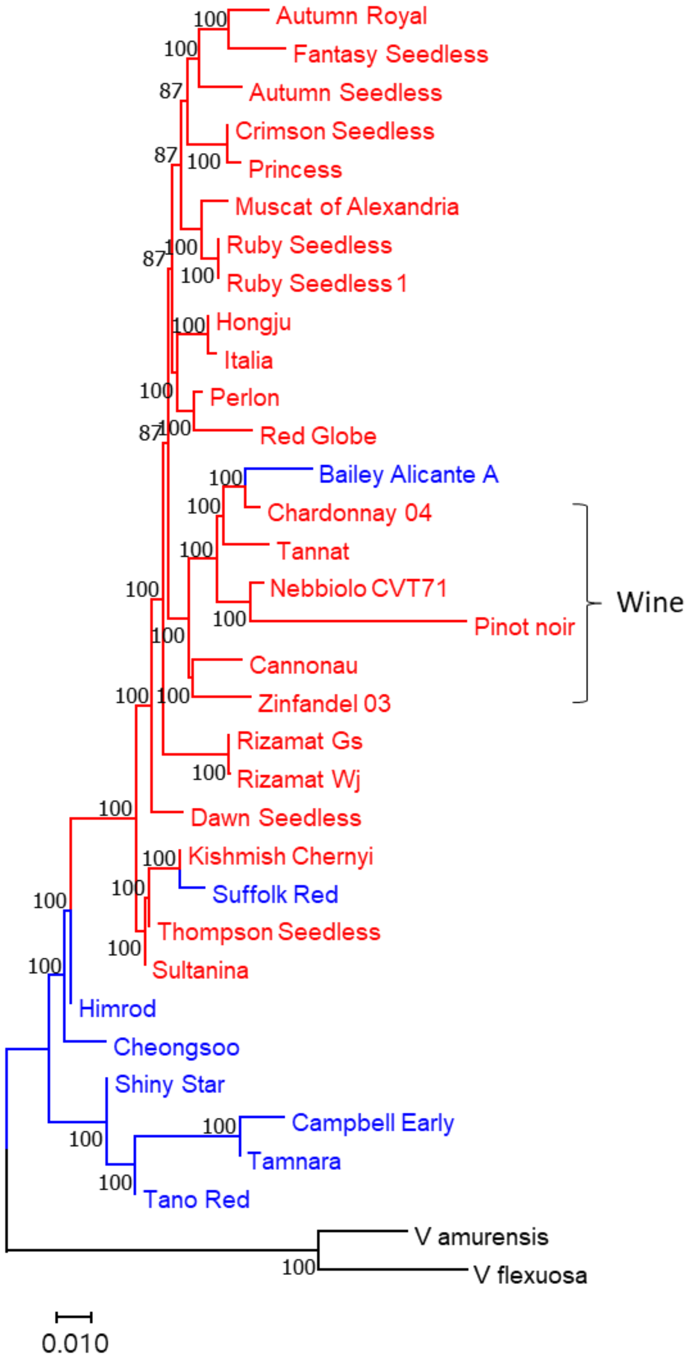


Figure S6. Neighbor-joining phylogenetic tree of 34 grape nuclear genomes constructed using the 17.45 million candidate SNP data without minor allele frequency filtration called from whole genome resequencing data. The evolutionary distances were measured by *p*-distance and branch supports were inferred with 1000 bootstrap replicates. Accessions in the neighbor-joining tree are represented by different colors: *Vitis vinifera* (red), interspecific hybrids (blue), and wild relatives (black). Group of wine grape accessions is indicated to emphasize their unique pedigree in this tree.

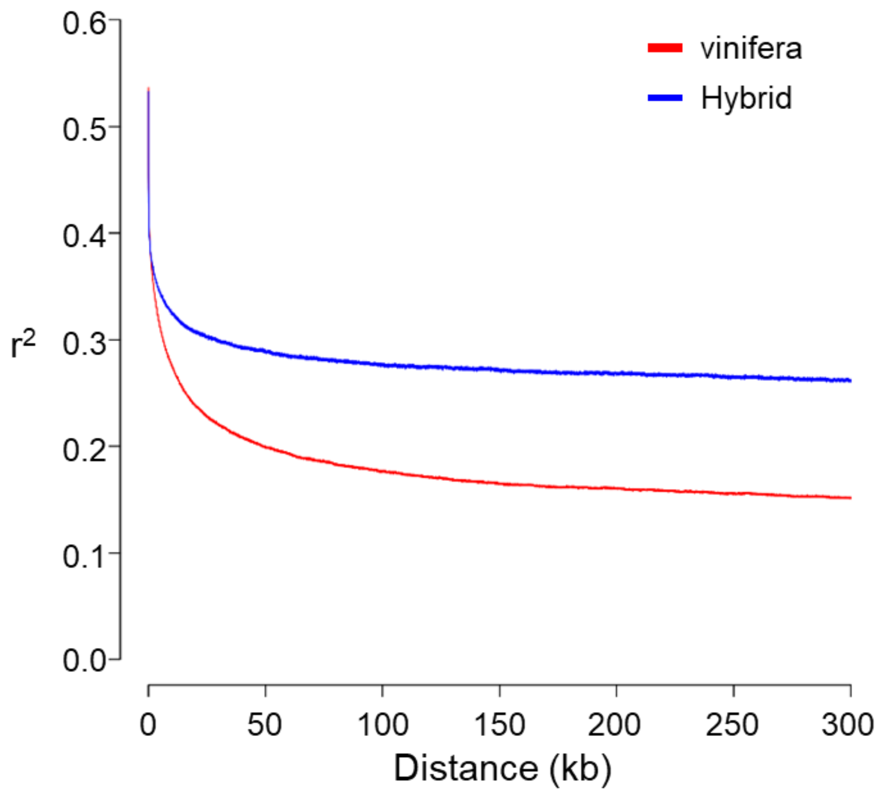


Figure S7. Curves of linkage disequilibrium decay patterns determined by squared correlations of allele frequencies ($r^2$) against distance between polymorphic sites in 20 *Vitis vinefera* (red) and eight interspecific hybrid (blue) grape accessions.

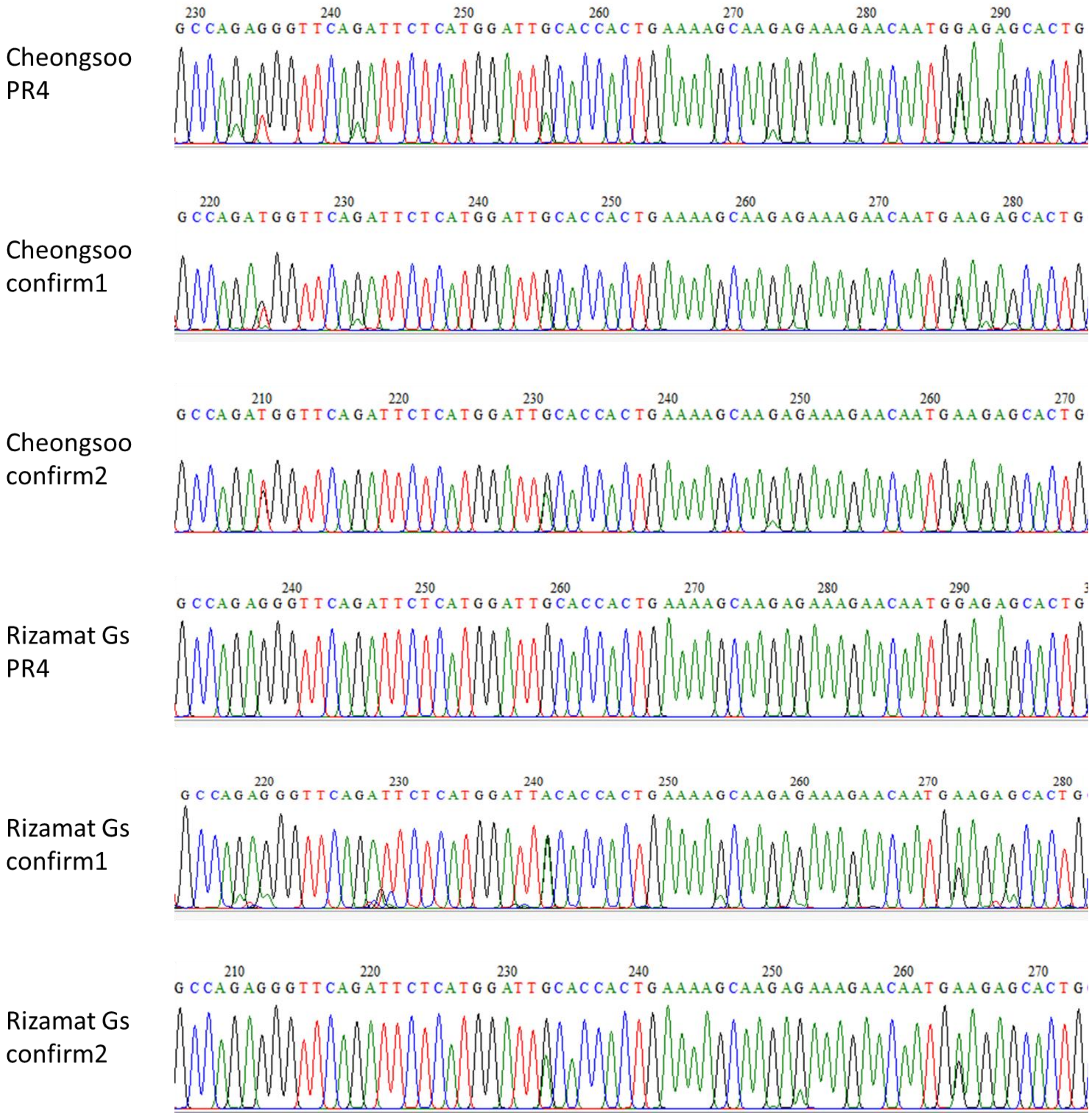


Figure S8. Representative Sanger sequencing profiles that show overlapping peaks PCR-amplified from two haplotype DNA fragments. PR4 are parts of profiles obtained by sequencing PCR products amplified from a pair of VviAGL11-PF2 and VviAGL11-PR4 primers using VviAGL11-PR4 primer. Confirm1 and confirm2 are parts of profiles obtained by sequencing approximately 1-kb sizes of PCR products amplified from two newly designed primer pairs, confirm1 and confirm2, using each of their forward primers. Rizamat Gs PR showed no overlapping peaks, however Rizamat Gs confirm1 and Rizamat Gs confirm2 showed overlapping peaks, indicating preferential PCR amplification of one of two haplotypes with the PCR amplification using the VviAGL11-PF2 and VviAGL11-PR4 primer pair.

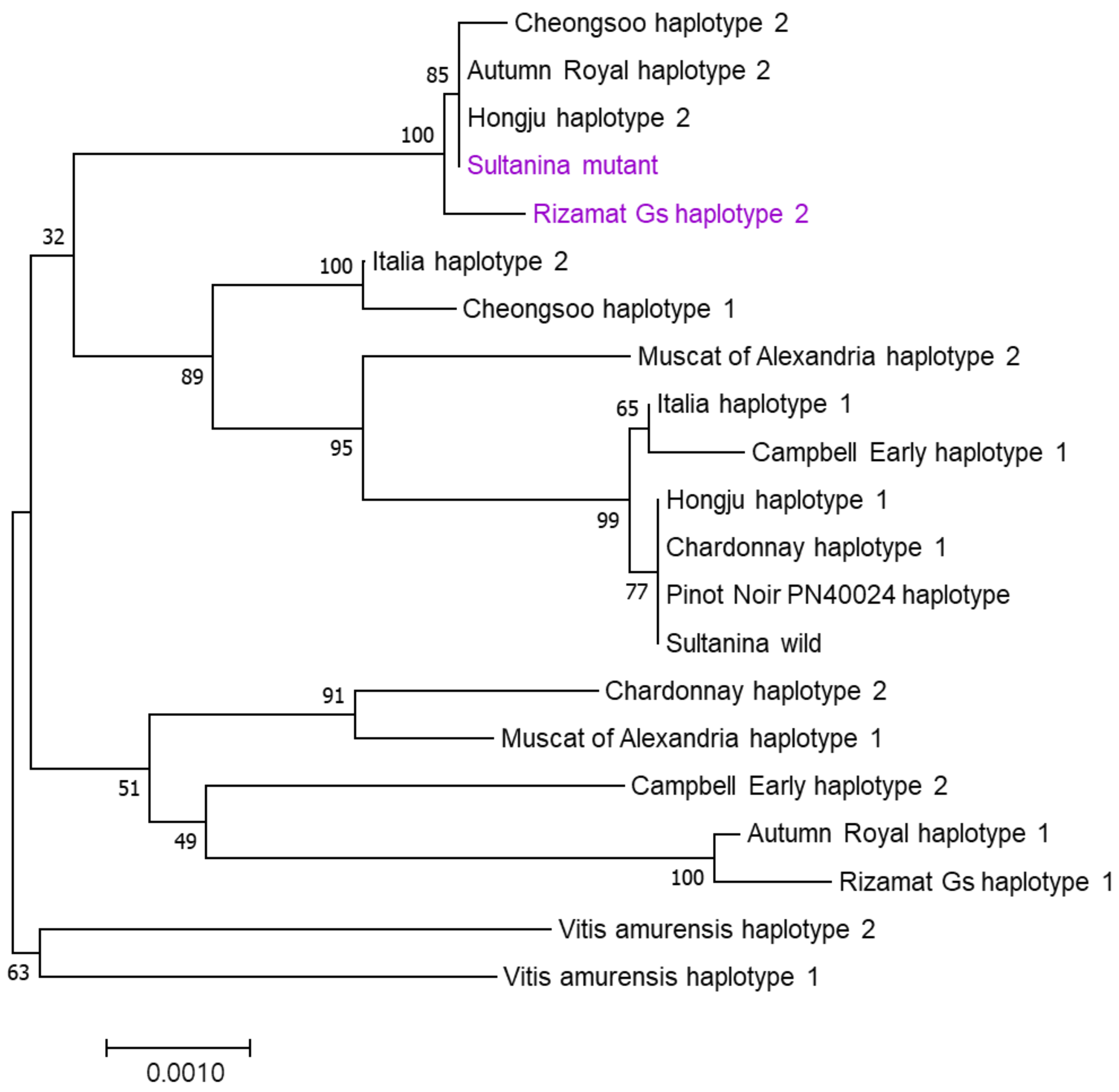


Figure S9. Neighbor-joining phylogenetic tree constructed from 3.1 kb-genomic DNA sequences including promoter and N-terminal coding regions of the *VviAGL11* gene. The evolutionary distances were measured by *p*-distance and branch supports were inferred with 1000 bootstrap replicates. Sultanina mutant and Rizamat Gs haplotype 2 are highlighted by purple letters.

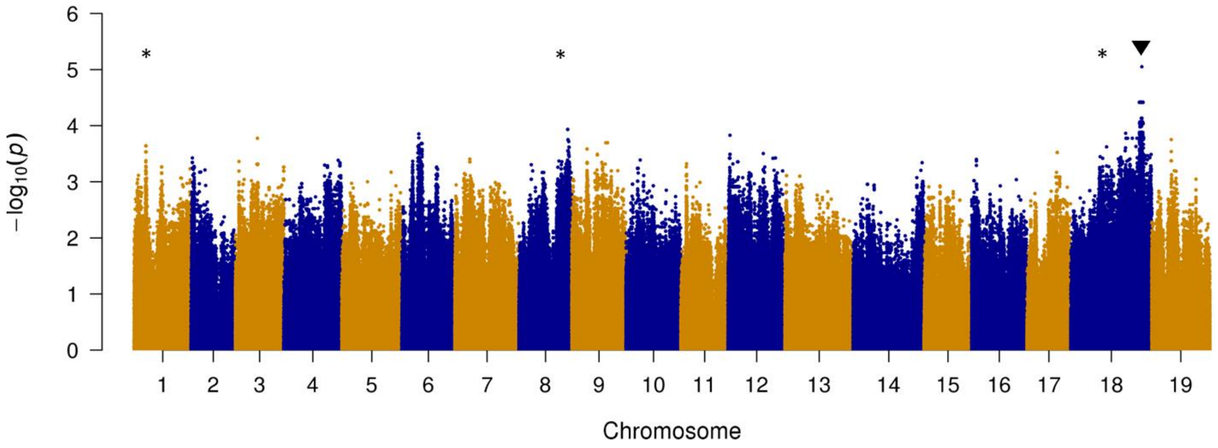


Figure S10. Manhattan plots of seedlessness genome-wide logistic association scans using GENESIS (Conomos et al., 2019). The -log10 *P* values from a genome-wide scan are plotted against the position on each of the 19 grape chromosomes. Chromosomal locations of *Vitvi01g00455*, *Vitvi08g01528*, and *Vitvi18g01237* associated with the minor seedless-regulating QTL and AGL11 associated with a major dominant QTL, which were predicted based on this GENESIS analysis as well as XP-CLR and SIFT (Sorting Intolerant From Tolerant) analyses, are indicated by * and ▼, respectively.

Table S1. Primers for sequencing used in this study

| Target gene | RefSeqID | Use | Name | Orientation | Sequence (5'-3') | Predicted PCR product size |
|---|---|---|---|---|---|---|
| AGL11 | LOC100232870 | PCR and sequencing | AGL11p | Forward | ATGCATTATTACCCTTCATA | 3770 |
| | | | | Reverse | CACCAAAACTCACATAAGAT | |
| | | PCR and sequencing | AGL11t | Forward | GCATGTTAATTAGTTTGACC | 3813 |
| | | | | Reverse | GTGATATATGGTGCTTCATT | |
| | | Sequencing | A11t-seqF1 | Forward | GGACAAGACTTTTCAACTGC | |
| | | | A11t-seqF2 | Forward | AATTGAGCTGGAAAATGAAA | |
| | | | A11t-seqR1 | Reverse | TGATAGCTCAATCAATTGTTAAAT | |
| | | | A11t-seqR1m | Reverse | TGATAACTCAATCAATTGTGGAA | |
| | | PCR and sequencing | A11-PF1 | Forward | CCACTGATATGGATTGATTTGCC | 3027 |
| | | | | Reverse | GGACAGTGGAATACAGATTC | |
| | | Sequencing | PF1-seqr1 | Reverse | AGGTTTATTGCACCAAAAGA | |
| | | PCR and sequencing | A11-PF2 | Forward | TAGGAAGGGATTACATGGG | 3552 |
| | | | A11-PR4 | Reverse | GAGTTTGTATGGAGAATAGCAG | |
| | | Sequencing | A11-PF3 | Forward | CCCATTACATCTTTGTGTGG | |
| | | | A11-PR3 | Reverse | AAACCCCACTGTGATAGG | |
| | | | A11-PF4 | Forward | GGATGGTGTCATAATGATAGG | |
| | | PCR and sequencing | confirm1 | Forward | TGGAGAATAGCAGTTGAAAAG | 909 |
| | | | | Reverse | TGAGAGCCTATTTGGGATAA | |
| | | PCR and sequencing | confirm2 | Forward | TGAAAAGTCTTGTCTTGGTTT | 900 |
| | | | | Reverse | AAAAATGAGAGCCTATTTGG | |
| AGL15 | LOC100253397 | PCR and sequencing | AGL15 | Forward | GGGTATCCAAATAACTCTCT | 1647 |
| | | | | Reverse | GTAACCAAACAGTCAAAAAG | |
| EXP-B3 | LOC100260198 | PCR and sequencing | expansin | Forward | TTTTACCAAAGTCAATGTTT | 1563 |

| | | | | | | |
|---|---|---|---|---|---|---|
| | | | | Reverse | TTGTCTTTCTTTGTTTGTCT | |
| FER1 | LOC100249810 | PCR and sequencing | FER_a | Forward | ATGATATTTTCCTTCCTCTC | 1582 |
| | | | | Reverse | TACATTTGCTTCCAGTAGAT | |
| FER2 | LOC100246444 | PCR and sequencing | FER_c | Forward | GTAAGTGGGACCTATTACCT | 1767 |
| | | | | Reverse | CATTGATGTAACCTTTGAAT | |
| GA2OX2 | LOC100244543 | PCR and sequencing | GA2ox2 | Forward | TAGGTTCCACTCTTTTATCA | 1855 |
| | | | | Reverse | AGGTGCTATGATTTCAGTTA | |
| MYB139 | LOC100246160 | PCR and sequencing | MYB139 | Forward | AAAAATAGCTGATAAAGCAA | 2386 |
| | | | | Reverse | GTCTGCAATAATGATAGAGC | |
| | | Sequencing | MYB139seqr | Reverse | TTCCTCCTTACTGAAGTTTCC | |
| MYB190 | LOC100259031 | PCR and sequencing | MYB190 | Forward | GTAAGTAGAATACGGACACG | 1534 |
| | | | | Reverse | TGGACCACCTAGTAATTAAA | |
| SND2 | LOC100267787 | PCR and sequencing | SND2 | Forward | ACTCAACCCTTTTTAATCTT | 2174 |
| | | | | Reverse | AGCATGCATTACTCATAACT | |
| | | Sequencing | SND2-seqf | Forward | TTTCTCACTTGTGTCATTGTTT | |
| SND3 | LOC100250388 | PCR and sequencing | SND3 | Forward | AATAAACCAAAACTCCTCTT | 1723 |
| | | | | Reverse | TAATGGAGGTCTTATTGAAA | |
| PIF4X2 | LOC100262490 | PCR and sequencing | PIF4 | Forward | TACTTGAATTCAGCTGTTCT | 1509 |
| | | | | Reverse | TTGAGTACAAGGATCATTTC | |
| THE1X4 | LOC100241800 | PCR and sequencing | THE1 | Forward | TCTTGTTGAGTTGAGTTTTT | 2425 |
| | | | | Reverse | ACATCAGATTTTTCTGTGAG | |
| | | Sequencing | THE1-seqf | Forward | CTTGGGAGAATGATGAGAAA | |
| | | | THE1-seqr | Reverse | ATTGGTGGAAGCTAATGAGA | |

Table S2. Percent concordance of candidate SNPs called through UnifiedGenotyper and HaplotyperCaller against SNPs called through Sanger sequencing data

| Species | Accession name | Sequenced gene region[a] | Total length of sequence (kb) | Number of candidate SNPs from UnifiedGenotyper | Percent concordance | | |
|---|---|---|---|---|---|---|---|
| | | | | | UnifiedGenotyper | Adjustment of heterozygous genotypes | HaplotyperCaller |
| *Vitis vinifera* | Autumn Royal | *AGL11, MYB190, SND2* | 6.7 | 104 | 93.3 | 92.1 | 94.5 |
| *Vitis vinifera* | Chardonnay | *AGL11* | 8.6 | 203 | 97.5 | 97.5 | 97.1 |
| *Vitis vinifera* | Hongju | *AGL11, EXP-B3, MYB190* | 9.8 | 204 | 93.1 | 92.4 | 94 |
| *Vitis vinifera* | Italia | *AGL11, GA2OX2, MYB190, SND2 , SND3, THE1X4* | 15.9 | 218 | 97.7 | 98.6 | 97.3 |
| *Vitis vinifera* | Muscat of Alexandria | *AGL11, FER2, MYB190, PIF4X2, THE1X4* | 9.9 | 100 | 94 | 93.9 | 95.3 |
| *Vitis vinifera* | Rizamat Gs | *AGL11, SND2* | 5.2 | 92 | 1 | 98.9 | 93.8 |
| *Vitis vinifera* | Sultanina | *AGL11* | 8.6 | 202 | 97 | 97.5 | 97.7 |
| **Total for *V. vinifera*** | | | **64.7** | **1123** | **96.2** | **96.1** | **96** |
| *Vitis labrusca* x *V. vinifera* | Campbell Early | *AGL11, SND2* | 9.0 | 220 | 75.9 | 97.7 | 28.7 |
| *Vitis* sp. | Cheongsoo | *AGL11, EXP-B3, MYB190, SND2, SND3* | 14.2 | 255 | 90.6 | 97.6 | 47 |
| **Total for hybrid** | | | **23.2** | **475** | **83.8** | **97.6** | **38.6** |
| **Vitus amurensis** | Vitus amurensis | *AGL11, FER2, GA2OX2, MYB190, SND2* | **10.3** | **102** | **87.3** | **92.9** | **55.8** |
| **Grand total** | | | **98.2** | **1700** | **92.2** | **96.2** | **77.4** |

[a] List of genes whose parts were sequenced.

Notes: We attempted to sequence approximately 36 kb of grape genomic DNA from twelve different genes from each of six selected accessions. SNPs called from Sanger sequences were compared against SNP calls from UnifiedGenotyper and HaplotypeCaller, respectively. Among these sequences, those from the *AGL15 and MYB139* genomic regions were excluded because we obtained more than three haplotypes from these regions, indicating that the grape genomes contain more than two paralogs of these genes. However, Blast searches of these against the grape reference genome sequence clearly indicated that the present reference contains only one copy of these sequences. This is likely because the paralog is absent in the grape reference genome. Sequences from a *FER1* genomic region were also excluded. Although *FER1*sequences showed one or two SNPs against the reference genome sequence, variant calling results showed unusually high numbers of heterozygous variations in both UnifiedGenotyper and HaplotypeCaller SNP calls. We obtained three haplotype sequences of *SND2* from Muscat of Alexandria, indicating that this gene has been duplicated in Muscat of Alexandria genome. Thus, *SND2* sequences from Muscat of Alexandria were excluded from this SNP concordance test. AGL11 sequences (GenBank Accession no. KM401845 - KM401848) of Chardonnay and Sultanina was previously reported by Malabarba et al. (2017).

Table S3. Number of homozygous SNPs with higher than 30% of allelic balance.

| Accession name | Group | 0/0 (reference homozygous) | 1/1 (non-reference homozygous) |
|---|---|---|---|
| Autumn Royal | *vinifera* | 1,389 | 2,432 |
| Autumn Seedless | *vinifera* | 1,144 | 1,959 |
| Cannonau | *vinifera* | 4,832 | 7,940 |
| Chardonnay 04 | *vinifera* | 1,934 | 3,055 |
| Crimson Seedless | *vinifera* | 2,304 | 3,799 |
| Dawn Seedless | *vinifera* | 1,345 | 1,982 |
| Fantasy Seedless | *vinifera* | 1,041 | 2,129 |
| Hongju | *vinifera* | 1,798 | 2,976 |
| Italia | *vinifera* | 854 | 1,344 |
| Kishmish Chernyi | *vinifera* | 4,740 | 7,971 |
| Muscat of Alexandria | *vinifera* | 7,256 | 11,254 |
| Nebbiolo_CVT71 | *vinifera* | 1,992 | 2,755 |
| Perlon | *vinifera* | 4,455 | 6,568 |
| Princess | *vinifera* | 8,058 | 12,076 |
| Red Globe | *vinifera* | 5,403 | 8,105 |
| Rizamat Gs | *vinifera* | 6,833 | 10,695 |
| Rizamat Wj | *vinifera* | 7,660 | 11,864 |
| Ruby Seedless | *vinifera* | 9,334 | 13,718 |
| Ruby Seedless 1 | *vinifera* | 9,470 | 14,610 |
| Sultanina | *vinifera* | 761 | 2,077 |
| Tannat | *vinifera* | 719 | 1,524 |
| Thompson Seedless | *vinifera* | 1,794 | 3,001 |
| Zinfandel 03 | *vinifera* | 2,384 | 3,943 |
| Bailey Alicante A | Hybrid | 94,754 | 183,293 |
| Campbell Early | Hybrid | 304,102 | 815,270 |
| Cheongsoo | Hybrid | 275,558 | 550,421 |
| Himrod | Hybrid | 110,268 | 248,222 |
| Shiny Star | Hybrid | 125,677 | 266,720 |
| Suffolk Red | Hybrid | 180,319 | 359,136 |
| Tamnara | Hybrid | 383,085 | 925,976 |
| Tano Red | Hybrid | 133,184 | 288,114 |
| Vitus amurensis | Wild | 141,538 | 480,749 |
| Vitus flexuosa | Wild | 311,871 | 1,044,017 |

Table S4. Distribution of genotypes of 17,453,275 of candidate SNPs in each of grape accessions. Non-reference homozygous genotype is indicated by 1/1, heterozygous by 0/1, missing by ./., and reference homozygous by 0/0.

| Accession name | Group | 1/1 | 0/1 | 0/1+1/1 | ./. | 0/0 |
|---|---|---|---|---|---|---|
| Autumn Royal | *vinifera* | 1,855,196 | 2,057,455 | 3,912,651 | 731,820 | 12,808,804 |
| Autumn Seedless | *vinifera* | 1,866,013 | 2,348,884 | 4,214,897 | 423,223 | 12,815,155 |
| Cannonau | *vinifera* | 999,575 | 1,900,844 | 2,900,419 | 5,291,134 | 9,261,722 |
| Chardonnay 04 | *vinifera* | 1,118,357 | 2,781,550 | 3,899,907 | 685,665 | 12,867,703 |
| Crimson Seedless | *vinifera* | 1,612,492 | 2,596,825 | 4,209,317 | 587,135 | 12,656,823 |
| Dawn Seedless | *vinifera* | 1,562,722 | 2,416,955 | 3,979,677 | 361,208 | 13,112,390 |
| Fantasy Seedless | *vinifera* | 2,068,229 | 1,978,254 | 4,046,483 | 610,481 | 12,796,311 |
| Hongju | *vinifera* | 1,544,258 | 2,538,163 | 4,082,421 | 646,601 | 12,724,253 |
| Italia | *vinifera* | 1,398,718 | 2,797,944 | 4,196,662 | 327,253 | 12,929,360 |
| Kishmish Chernyi | *vinifera* | 1,771,808 | 2,582,379 | 4,354,187 | 558,323 | 12,540,765 |
| Muscat of Alexandria | *vinifera* | 1,555,577 | 2,676,247 | 4,231,824 | 569,361 | 12,652,090 |
| Nebbiolo_CVT71 | *vinifera* | 988,576 | 2,937,421 | 3,925,997 | 600,656 | 12,926,622 |
| Perlon | *vinifera* | 1,566,318 | 2,725,533 | 4,291,851 | 414,313 | 12,747,111 |
| Princess | *vinifera* | 1,914,816 | 2,219,396 | 4,134,212 | 368,804 | 12,950,259 |
| Red Globe | *vinifera* | 1,761,902 | 2,259,824 | 4,021,726 | 398,405 | 13,033,144 |
| Rizamat Gs | *vinifera* | 1,659,837 | 2,624,508 | 4,284,345 | 345,005 | 12,823,925 |
| Rizamat Wj | *vinifera* | 1,662,339 | 2,610,950 | 4,273,289 | 350,490 | 12,829,496 |
| Ruby Seedless | *vinifera* | 1,602,615 | 2,708,613 | 4,311,228 | 305,233 | 12,836,814 |
| Ruby Seedless 1 | *vinifera* | 1,585,802 | 2,675,070 | 4,260,872 | 429,580 | 12,762,823 |
| Sultanina | *vinifera* | 1,598,696 | 2,761,874 | 4,360,570 | 495,782 | 12,596,923 |
| Tannat | *vinifera* | 1,319,509 | 2,886,057 | 4,205,566 | 374,923 | 12,872,786 |
| Thompson Seedless | *vinifera* | 1,668,236 | 2,825,301 | 4,493,537 | 308,759 | 12,650,979 |
| Zinfandel 03 | *vinifera* | 1,537,454 | 2,690,543 | 4,227,997 | 763,645 | 12,461,633 |
| Bailey Alicante A | Hybrid | 1,463,722 | 3,400,202 | 4,863,924 | 759,574 | 11,829,777 |
| Campbell Early | Hybrid | 2,148,416 | 2,783,137 | 4,931,553 | 2,683,117 | 9,838,605 |
| Cheongsoo | Hybrid | 1,838,860 | 3,552,195 | 5,391,055 | 917,010 | 11,145,210 |
| Himrod | Hybrid | 1,644,539 | 3,835,841 | 5,480,380 | 911,830 | 11,061,065 |
| Shiny Star | Hybrid | 1,906,860 | 3,245,484 | 5,152,344 | 1,361,054 | 10,939,877 |
| Suffolk Red | Hybrid | 1,549,519 | 3,417,627 | 4,967,146 | 2,361,113 | 10,125,016 |
| Tamnara | Hybrid | 2,225,163 | 3,046,121 | 5,271,284 | 2,098,720 | 10,083,271 |
| Tano Red | Hybrid | 1,766,698 | 3,463,227 | 5,229,925 | 924,470 | 11,298,880 |
| Vitus amurensis | Wild | 2,747,907 | 1,790,225 | 4,538,132 | 3,534,615 | 9,380,528 |
| Vitus flexuosa | Wild | 2,159,522 | 3,291,327 | 5,450,849 | 5,867,153 | 6,135,273 |

Table S5. Distribution of genotypes of 5,373,452 of candidate SNPs in each of grape accessions. Non-reference homozygous genotype is indicated by 1/1, heterozygous by 0/1, missing by ./., and reference homozygous by 0/0.

| Accession name | Class | 1/1 | 0/1 | 0/1+1/1 | ./. | 0/0 |
|---|---|---|---|---|---|---|
| Autumn Royal | *vinifera* | 1,047,726 | 1,340,631 | 2,388,357 | 75,233 | 2,909,862 |
| Autumn Seedless | *vinifera* | 1,022,569 | 1,545,523 | 2,568,092 | 38,782 | 2,766,578 |
| Cannonau | *vinifera* | 569,611 | 1,193,461 | 1,763,072 | 1,444,187 | 2,166,193 |
| Chardonnay 04 | *vinifera* | 597,976 | 1,619,525 | 2,217,501 | 95,442 | 3,060,509 |
| Crimson Seedless | *vinifera* | 851,076 | 1,746,014 | 2,597,090 | 65,001 | 2,711,361 |
| Dawn Seedless | *vinifera* | 849,038 | 1,640,820 | 2,489,858 | 37,350 | 2,846,244 |
| Fantasy Seedless | *vinifera* | 1,180,886 | 1,254,879 | 2,435,765 | 53,441 | 2,884,246 |
| Hongju | *vinifera* | 825,400 | 1,694,642 | 2,520,042 | 71,761 | 2,781,649 |
| Italia | *vinifera* | 722,256 | 1,754,012 | 2,476,268 | 29,587 | 2,867,597 |
| Kishmish Chernyi | *vinifera* | 984,083 | 1,630,677 | 2,614,760 | 54,745 | 2,703,947 |
| Muscat of Alexandria | *vinifera* | 824,789 | 1,674,966 | 2,499,755 | 64,414 | 2,809,283 |
| Nebbiolo_CVT71 | *vinifera* | 516,925 | 1,646,649 | 2,163,574 | 89,867 | 3,120,011 |
| Perlon | *vinifera* | 812,448 | 1,848,691 | 2,661,139 | 41,309 | 2,671,004 |
| Princess | *vinifera* | 1,066,346 | 1,487,371 | 2,553,717 | 29,744 | 2,789,991 |
| Red Globe | *vinifera* | 938,905 | 1,459,795 | 2,398,700 | 35,648 | 2,939,104 |
| Rizamat Gs | *vinifera* | 891,993 | 1,619,701 | 2,511,694 | 32,905 | 2,828,853 |
| Rizamat Wj | *vinifera* | 893,422 | 1,615,412 | 2,508,834 | 34,413 | 2,830,205 |
| Ruby Seedless | *vinifera* | 815,986 | 1,834,985 | 2,650,971 | 28,080 | 2,694,401 |
| Ruby Seedless 1 | *vinifera* | 815,287 | 1,818,823 | 2,634,110 | 47,891 | 2,691,451 |
| Sultanina | *vinifera* | 848,589 | 1,950,888 | 2,799,477 | 81,193 | 2,492,782 |
| Tannat | *vinifera* | 685,443 | 1,598,079 | 2,283,522 | 41,226 | 3,048,704 |
| Thompson Seedless | *vinifera* | 869,185 | 1,974,936 | 2,844,121 | 31,248 | 2,498,083 |
| Zinfandel 03 | *vinifera* | 839,283 | 1,580,619 | 2,419,902 | 100,050 | 2,853,500 |
| Bailey Alicante A | Hybrid | 738,094 | 1,764,578 | 2,502,672 | 92,234 | 2,778,546 |
| Campbell Early | Hybrid | 1,358,617 | 1,413,245 | 2,771,862 | 336,682 | 2,264,908 |
| Cheongsoo | Hybrid | 1,003,900 | 2,077,013 | 3,080,913 | 87,369 | 2,205,170 |
| Himrod | Hybrid | 853,788 | 2,515,089 | 3,368,877 | 95,082 | 1,909,493 |
| Shiny Star | Hybrid | 1,102,159 | 2,093,932 | 3,196,091 | 128,165 | 2,049,196 |
| Suffolk Red | Hybrid | 868,846 | 2,106,924 | 2,975,770 | 388,670 | 2,009,012 |
| Tamnara | Hybrid | 1,349,613 | 1,704,528 | 3,054,141 | 240,558 | 2,078,753 |
| Tano Red | Hybrid | 952,945 | 2,111,979 | 3,064,924 | 82,267 | 2,226,261 |
| Vitus amurensis | Wild | 1,352,919 | 431,168 | 1,784,087 | 765,599 | 2,823,766 |
| Vitus flexuosa | Wild | 1,032,179 | 894,900 | 1,927,079 | 1,494,989 | 1,951,384 |

Table S6. List of chromosomal locations of the highest XP-CLR points in 30 candidate selective sweeps detected

| Chromosome | Range of 5-kb window | | |
|---|---|---|---|
| | Start | End | XP-CLR value |
| 1 | 4,882,001 | 4,887,000 | 13.57513 |
| 1 | 14,734,001 | 14,739,000 | 9.413687 |
| 1 | 21,797,001 | 21,802,000 | 11.34642 |
| 3 | 9,261,001 | 9,266,000 | 19.83749 |
| 5 | 16,613,001 | 16,618,000 | 10.28197 |
| 6 | 7,666,001 | 7,671,000 | 17.61092 |
| 6 | 13,241,001 | 13,246,000 | 11.90963 |
| 7 | 6,839,001 | 6,844,000 | 34.76061 |
| 7 | 11,711,001 | 11,716,000 | 10.01538 |
| 8 | 9,691,001 | 9,696,000 | 21.02209 |
| 8 | 17,975,001 | 17,980,000 | 32.8218 |
| 8 | 20,701,001 | 20,706,000 | 30.5868 |
| 9 | 6,440,001 | 6,445,000 | 15.15622 |
| 9 | 13,984,001 | 13,989,000 | 16.42361 |
| 9 | 16,022,001 | 16,027,000 | 29.03137 |
| 10 | 4,842,001 | 4,847,000 | 10.4536 |
| 11 | 2,413,001 | 2,418,000 | 9.855065 |
| 11 | 7,307,001 | 7,312,000 | 14.27136 |
| 11 | 11,702,001 | 11,707,000 | 11.9245 |
| 12 | 703,001 | 708,000 | 13.02817 |
| 13 | 8,772,001 | 8,777,000 | 10.8748 |
| 13 | 21,288,001 | 21,293,000 | 16.36479 |
| 14 | 29,382,001 | 29,387,000 | 33.73349 |
| 15 | 3,230,001 | 3,235,000 | 18.17089 |
| 16 | 21,121,001 | 21,126,000 | 28.777 |
| 18 | 8,970,001 | 8,975,000 | 13.4437 |
| 18 | 13,336,001 | 13,341,000 | 20.75844 |
| 18 | 26,107,001 | 26,112,000 | 23.5555 |
| 18 | 29,962,001 | 29,967,000 | 28.15971 |
| 19 | 10,129,001 | 10,134,000 | 8.597111 |

Table S7. List of 41 deleterious SNPs with -log10 P value of > 2.5 from GWAS and distribution of their genotypes in 15 seedless and 13 seeded grape accessions.

| Chromosome | Position | -log *P* | Gene ID | Accession number in each of genotypes[1] | | | | Accession number in each of genotypes | | | |
|---|---|---|---|---|---|---|---|---|---|---|---|
| | | | | 0/0 | 0/1 | 1/1 | ./. | 0/0 | 0/1 | 1/1 | ./. |
| 1 | 4908460 | 2.909 | Vitvi01g00447 | 0 | 4 | 10 | 0 | 7 | 5 | 1 | 0 |
| 1 | 4911453 | 2.909 | Vitvi01g00447 | 0 | 3 | 10 | 1 | 7 | 5 | 1 | 0 |
| 1 | 4912938 | 2.909 | Vitvi01g00448 | 0 | 4 | 10 | 0 | 7 | 5 | 1 | 0 |
| 1 | 4913189 | 2.909 | Vitvi01g00448 | 0 | 4 | 10 | 0 | 7 | 5 | 1 | 0 |
| 1 | 4973655 | 2.909 | Vitvi01g00453 | 0 | 4 | 10 | 0 | 6 | 5 | 1 | 1 |
| 1 | 5006281 | 3.05 | Vitvi01g00455 | 0 | 4 | 10 | 0 | 9 | 4 | 0 | 0 |
| 3 | 9396594 | 2.553 | Vitvi03g00780 | 3 | 10 | 2 | 0 | 12 | 1 | 0 | 0 |
| 6 | 7513179 | 2.733 | Vitvi06g00662 | 2 | 6 | 7 | 0 | 11 | 2 | 0 | 0 |
| 6 | 7757492 | 2.895 | Vitvi06g00687 | 3 | 8 | 4 | 0 | 13 | 0 | 0 | 0 |
| 6 | 8146919 | 2.552 | Vitvi06g00725 | 2 | 5 | 8 | 0 | 9 | 4 | 0 | 0 |
| 7 | 6342363 | 3.355 | Vitvi07g00586 | 1 | 5 | 8 | 0 | 10 | 3 | 0 | 0 |
| 7 | 6867830 | 2.867 | Vitvi07g00618 | 2 | 9 | 4 | 0 | 12 | 1 | 0 | 0 |
| 7 | 6896568 | 2.867 | Vitvi07g00620 | 2 | 9 | 4 | 0 | 12 | 1 | 0 | 0 |
| 7 | 6949284 | 2.576 | Vitvi07g00624 | 2 | 9 | 4 | 0 | 11 | 2 | 0 | 0 |
| 7 | 6950935 | 2.576 | Vitvi07g00624 | 2 | 8 | 4 | 1 | 11 | 2 | 0 | 0 |
| 7 | 7083128 | 2.576 | Vitvi07g00633 | 2 | 8 | 4 | 1 | 11 | 2 | 0 | 0 |
| 7 | 7088939 | 2.576 | Vitvi07g00633 | 2 | 9 | 4 | 0 | 11 | 2 | 0 | 0 |
| 8 | 17965153 | 3.104 | Vitvi08g01518 | 1 | 7 | 7 | 0 | 11 | 2 | 0 | 0 |
| 8 | 18038427 | 2.512 | Vitvi08g01528 | 0 | 3 | 11 | 0 | 7 | 4 | 2 | 0 |
| 8 | 20261534 | 2.922 | Vitvi08g01741 | 1 | 12 | 2 | 0 | 11 | 2 | 0 | 0 |

| | | | | | | | | | | | |
|---|---|---|---|---|---|---|---|---|---|---|---|
| 8 | 20734925 | 3.343 | Vitvi08g02366 | 1 | 11 | 2 | 1 | 12 | 1 | 0 | 0 |
| 8 | 20806690 | 3.343 | Vitvi08g01790 | 1 | 12 | 2 | 0 | 12 | 1 | 0 | 0 |
| 8 | 20982004 | 2.668 | Vitvi08g01817 | 4 | 8 | 2 | 0 | 13 | 0 | 0 | 0 |
| 8 | 21011730 | 2.668 | Vitvi08g01825 | 4 | 8 | 2 | 0 | 13 | 0 | 0 | 0 |
| 8 | 21127209 | 2.517 | Vitvi08g02370 | 0 | 3 | 12 | 0 | 5 | 5 | 2 | 1 |
| 8 | 21127212 | 3.12 | Vitvi08g02370 | 0 | 4 | 11 | 1 | 7 | 5 | 0 | 1 |
| 8 | 21127278 | 2.959 | Vitvi08g02370 | 0 | 5 | 11 | 1 | 7 | 6 | 0 | 0 |
| 8 | 21185902 | 2.668 | Vitvi08g01847 | 4 | 9 | 2 | 0 | 13 | 0 | 0 | 0 |
| 9 | 6144974 | 2.556 | Vitvi09g01650 | 0 | 10 | 3 | 2 | 10 | 3 | 0 | 0 |
| 12 | 1098595 | 2.672 | Vitvi12g00059 | 5 | 10 | 0 | 0 | 13 | 0 | 0 | 0 |
| 14 | 29689361 | 3.003 | Vitvi14g01983 | 0 | 7 | 8 | 0 | 8 | 5 | 0 | 0 |
| 18 | 13614293 | 2.527 | Vitvi18g01230 | 0 | 5 | 10 | 0 | 4 | 7 | 2 | 0 |
| 18 | 13663964 | 2.814 | Vitvi18g01237 | 0 | 3 | 11 | 0 | 3 | 8 | 2 | 0 |
| 18 | 13738152 | 2.599 | Vitvi18g01245 | 0 | 4 | 10 | 1 | 4 | 8 | 1 | 0 |
| 18 | 29483428 | 4.416 | Vitvi18g02081 | 0 | 15 | 0 | 0 | 12 | 1 | 0 | 0 |
| 18 | 29983966 | 3.562 | Vitvi18g02110 | 0 | 13 | 2 | 0 | 10 | 1 | 0 | 2 |
| 18 | 30100746 | 3.341 | Vitvi18g02121 | 2 | 13 | 0 | 0 | 12 | 1 | 0 | 0 |
| 18 | 30103518 | 3.815 | Vitvi18g02121 | 1 | 13 | 0 | 1 | 12 | 1 | 0 | 0 |
| 18 | 30119603 | 3.753 | Vitvi18g02122 | 0 | 12 | 3 | 0 | 11 | 1 | 0 | 1 |
| 18 | 30306458 | 5.051 | Vitvi18g02133 | 0 | 15 | 0 | 0 | 12 | 0 | 0 | 1 |
| 18 | 30428349 | 3.316 | Vitvi18g02135 | 0 | 14 | 0 | 1 | 10 | 3 | 0 | 0 |

[1] 0/0 is reference homozygous, 0/1 heterozygous, 1/1 non-reference homozygous, ./. missing genotype.
